# Combined influence of food availability and agricultural intensification on a declining aerial insectivore

**DOI:** 10.1101/2021.02.02.427782

**Authors:** Daniel R. Garrett, Fanie Pelletier, Dany Garant, Marc Bélisle

## Abstract

Aerial insectivores show worldwide population declines coinciding with shifts in agricultural practices. Increasing reliance on certain agricultural practices is thought to have led to an overall reduction in insect abundance that negatively affects aerial insectivore fitness. The relationship between prey availability and the fitness of insectivores may thus vary with the extent of agricultural intensity. It is therefore imperative to quantify the strength and direction of these associations. Here we used data from an 11-year study monitoring the breeding of Tree Swallows (*Tachycineta bicolor*) and the availability of Diptera (their main prey) across a gradient of agricultural intensification in southern Québec, Canada. This gradient was characterized by a shift in agricultural production, whereby landscapes composed of forage and pastures represented less agro-intensive landscapes and those focusing on large-scale arable row crop monocultures, such as corn (*Zea mays*) or soybean (*Glycine max*) that are innately associated with significant mechanization and agro-chemical inputs, represented more agro-intensive landscapes. We evaluated the landscape characteristics affecting prey availability, and how this relationship influences the fledging success, duration of the nestling period, fledgling body mass, and wing length as these variables are known to influence the population dynamics of this species. Diptera availability was greatest within predominately forested landscapes, while within landscapes dominated by agriculture, it was marginally greater in less agro-intensive areas. Of the measured fitness and body condition proxies, both fledging success and nestling body mass were positively related to prey availability. The impact of prey availability varied across the agricultural gradient as fledging success improved with increasing prey levels within forage landscapes yet declined in more agro-intensive landscapes. Finally, after accounting for prey availability, fledging success was lowest, nestling periods were the longest, and wing length of fledglings were shortest within more agro-intensive landscapes. Our results highlight the interacting roles that aerial insect availability and agricultural intensification have on the fitness of aerial insectivores, and by extension how food availability may interact with other aspects of breeding habitats to influence the population dynamics of predators.

**Open Research:** Data are not yet provided (option 4) as they are being used for other research projects. We affirm that data will be permanently archived if the paper is accepted for publication on the Dryad repository.

## Introduction

Food availability is a key determinant of population size and dynamics (Hailman and Lack 1955, McCamey and Slobodkin 1963, Newton 1998). The amount of food resources an individual acquires from a given foraging effort affects the amount of energy it can allocate to both its survival and reproduction, thereby affecting vital rates (Martin 1987, Newton 1998, 2002, Stephens et al. 2007). Food reduction or supplementation experiments are often used to quantify the impact of food availability on reproductive output (Martin 1987, Seward et al. 2014, Ruffino et al. 2014, Seress et al. 2020). For example, *Bacillus thuringiensis* (Bt) insecticides are used to reduce insect abundance and both experimental and observational studies typically report reduced fitness of insectivorous predators following their application (Rodenhouse and Holmes 1992, Nagy and Smith 1997, Marshall et al. 2002, Poulin et al. 2010). Other studies, conducted under unaltered levels of food availability have reported no link with fitness outcomes of consumers (Dawson and Bortolotti 2000, Imlay et al. 2017, McClenaghan et al. 2019). Such contrasting results can be explained by food being overly abundant and thus other environmental (e.g., extreme temperature or elevated precipitation) or endogenous factors (e.g., parental condition, care, and past experience), having a greater influence than food availability in limiting reproductive output and subsequent population growth (Martin 1987, Newton 1998, White 2008, Winkler et al. 2013, Quinney 2019). Studies of naturally occurring predator-prey interactions have frequently been conducted within single landscapes, ones already suitable for elevated fitness, or have been conducted without reference to human dominated landscapes (Martin 1987, Fazey et al. 2005, Robb et al. 2008, Martin et al. 2012). Habitat induced variability in prey from such studies may thus not be great enough to influence the fitness of predators.

Human-driven landscape modifications are known to affect both the magnitude and direction of a broad spectrum of ecological processes occurring over a multitude of spatio-temporal scales (Rolstad 1991, Miller and Adams 1995, Fischer and Lindenmayer 2007, Davis 2007). Therefore, studies disregarding such impacts may overlook confounding or interacting factors of foraging within anthropogenic landscapes. For instance, populations of pest insects may be higher in landscapes with an abundance of large scale monocultures (Andow 1983, 1991). Yet in these same landscapes, the availability of other crucial resources may be limited, possibly influencing either the lifetime or annual fitness of animals continuing to perceive them as suitable breeding or foraging habitats (Robertson 2012, Hale et al. 2015, Letten et al. 2017). The historical removal of forests and replacement of grasslands and wetlands with agricultural lands, including the simplification of agricultural landscapes and a shift from mixed crop farming to livestock or highly-mechanized row-cropping supported by significant agro-chemical inputs, a process known as agricultural intensification, is one of the most prominent driver of human-induced landscape change (Lambin et al. 2001, Fischer and Lindenmayer 2007). In North America, this process is exemplified by the trend of agricultural landscapes transitioning from small scale perennial crops and low impact farming practices, to being increasingly composed of only a few arable row crop monocultures (e.g., corn (*Zea mays*) or soybean (*Glycine max*)), and the near ubiquitous use of the aforementioned agro-intensive practices (Stanton et al. 2018).

Despite agricultural intensification being linked to the decline of many vertebrates (Ceballos 2002, Rosenberg et al. 2019), the relationship between observed prey availability and the fitness of predators has rarely been investigated with respect to agricultural cover. For example, several species of grassland and farmland birds have displayed significant declines since the 1970s (Vickery et al. 2004, Boatman et al. 2004, Stanton et al. 2018, Rosenberg et al. 2019). In particular, steep declines are observed in many aerial insectivores, a guild of birds foraging nearly exclusively on aerial insects (Nebel et al. 2010, Michel et al. 2016, NABCI 2019). Due in part to the spatio-temporal correlations between these phenomena, a common hypothesis is that aerial insectivore declines are due in part to agricultural intensification influencing the abundance and species composition of the insects consumed by these birds (Stanton et al. 2018, Twining et al. 2018, Spiller and Dettmers 2019). Multiple lines of evidence exist linking changes in specific insect communities, abundance, behavior or phenology to agricultural intensification (Benton et al. 2002, Thomas 2004, Pisa et al. 2015, Stanton et al. 2018, Montgomery et al. 2020). Yet, the link between agricultural intensification and aerial insectivore declines, through a reduction in the availability of food resources, has so far yielded inconsistent results (Evans et al. 2007, Nocera et al. 2012, Imlay et al. 2017, Stanton et al. 2017, McClenaghan et al. 2019).

Within agricultural landscapes, practices such as large-scale, simplified arable row-cropping, increased reliance on agro-chemical inputs (i.e., pesticides and fertilizers) as well as the removal of both natural and marginal habitats are suspected to be the primary drivers of changes of insect populations (Tscharntke et al. 2005, Grüebler et al. 2008, Attwood et al. 2008, Montgomery et al. 2020). Furthermore, within heavily cultivated landscapes, the same mechanisms reducing insect abundance, may also impact insectivorous birds, either irrespective of, or in conjunction with, levels of prey. For example, due to the increase of pesticides, insectivorous birds breeding within landscapes dominated by agro-intensive monocultures may not only be exposed to decreased prey availability, but also to elevated toxicological loads of food items (Morrissey et al. 2015, Montiel-León et al. 2019, Malaj et al. 2020, Poisson et al. 2021). Breeding within agro-intensive landscapes may thus have the potential to impact fitness not only through a trophic pathway, via variation in prey availability, but also through other indirect pathways linked to agricultural intensification (Stanton et al. 2018).

Here, we present results of an 11-year study monitoring both the reproductive attempts of Tree Swallows (*Tachycineta bicolor*) and the availability of their aerial insect prey throughout a nest box network located along a gradient of agricultural intensification in southern Québec, Canada. Tree Swallows breed over much of North America and demonstrate population declines similar to those of most other aerial insectivores, especially in the north eastern portion of their breeding range (Shutler et al. 2012, Michel et al. 2016). Our principal objectives were to assess evidence with respect to three hypotheses. The first was that a gradient of agricultural intensification influences the availability of aerial insectivore prey, and we predicted that landscapes with increasing levels of arable row crops would result in a reduction in prey availability (Benton et al. 2002, Thomas 2004, Pisa et al. 2015, Stanton et al. 2018, Montgomery et al. 2020). The second hypothesis was that, as aerial insectivores, breeding Tree Swallows are food limited (Twining et al. 2018, Spiller and Dettmers 2019). Therefore, increased prey availability should result in greater fitness, including the number of nestlings surviving until fledging, reduced nestling duration (due to increased growth rates), and elevated nestling condition. Our third hypothesis was that Tree Swallow fitness is influenced by agricultural landcover through more than just the overall abundance of food, for example through pesticide exposure (Poisson et al. 2021) or variation in diet quality (Stanton et al. 2018, Twining et al. 2018). We therefore predicted that, after controlling for prey availability, breeding attempts within landscapes occupied by increasing row crops should result in decreased fitness.

Population dynamics of short lived species are thought to be driven by adult and juvenile return rates and to a lesser extent fecundity (Sæther and Bakke 2000). Within Tree Swallow populations, adult survival appears to be relatively stable throughout their distribution (Weegman et al. 2017, Clark et al. 2018). Furthermore, recent evidence showed that, due to their high elasticity, population viability of this species could largely be determined by fledgling survival and recruitment rates (Jones et al. 2017, Weegman et al. 2017, Cox et al. 2018). Although food limitation can be influential throughout the life cycle (Hostetler et al. 2015, Knight et al. 2019), given the above, we focus primarily on the nestling period, because factors influencing fledgling number and post fledging survival have a significant impact on population growth (Cox et al. 2018). We therefore assess the influence of prey availability on the fledging success, the duration of the nestling period, and the morphological traits of fledglings (hitherto referred collectively as fitness and body condition proxies). Evidence suggests food availability and agricultural landscapes can be drivers of both nestling growth and condition, and yet they have received less attention than annual reproductive output as a possible driver of aerial insectivore declines (Smith and Bruun 1998, Granbom and Smith 2006, Pigeon et al. 2013, Almasi et al. 2015, Kusack et al. 2020, Houle et al. 2020). Therefore, we assess the possibility that links between agricultural intensification and declining populations of aerial insectivores are not only through a reduction in the number of fledglings, but also through reduced offspring condition, as this may carry over and affect future recruitment and productivity (Stutchbury et al. 2011, Naef-Daenzer and Grüebler 2016, Saino et al. 2018, Evans et al. 2020, Jones and Ward 2020).

## Methods

### Study area and nest box system

Between 2006 and 2016, we monitored breeding attempts of Tree Swallows within 400 nest boxes dispersed throughout a gradient of agricultural intensification in southern Quebec, Canada.

Ten nest boxes were spaced approximately 50 m apart along the field margins of 40 different farms, located in various agricultural landscape contexts (Figure 1; see Ghilain and Bélisle 2008 for details). The gradient of agricultural intensification was characterized by an east-west shift of agricultural production. While the eastern portion of the system was composed primarily of pastures and forage crops (e.g., hay, alfalfa (*Medicago sativa*), and clover (*Trifolium spp.*)) interspersed with large expanses of forest cover, the west was dominated by large-scale monocultures (principally corn, soybean, and wheat (*Triticum spp.*)) and was denuded of forest cover (Bélanger and Grenier 2002, Jobin et al. 2005, Ruiz and Domon 2009). This increased use of monocultures has resulted in a near reliance on fertilizers, pesticides, mechanization and drainage of surface waters and wetlands (Jobin et al. 2003). Furthermore, between 2011 to 2019, approximately 100% of the corn and 60% of the soybean were sown as neonicotinoid-coated seeds in our study area (MDDELCC 2015). Subsequently, within the water bodies of the western part of our study area, neonicotinoids, as well as many other pesticides, are frequently detected at levels threatening to aquatic life if chronically exposed (Giroux 2019, Montiel-León et al. 2019).

**Figure 1:**
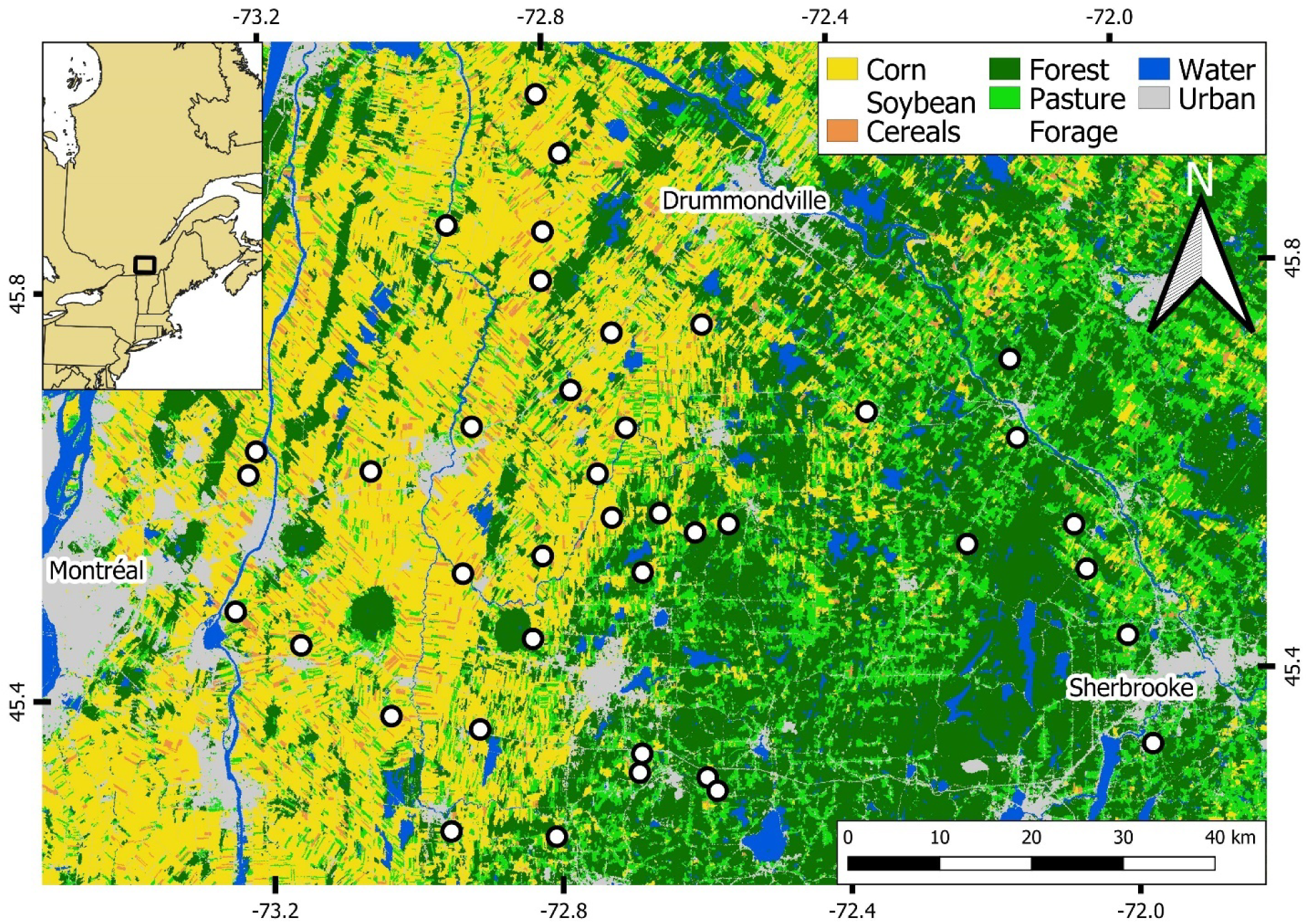
Map of the 40 different farms across a gradient of agricultural intensification within southern Québec, Canada. Each point represents the location of a farm. Underlying raster represents the agricultural gradient derived from the Annual crop inventory of 2013 (AAFC 2018), for which each pixel represents either one of the five higher order categories or open water. The rectangle within inset map represents the extent of the main map and thus the spatial extent of the study system. Graticules are latitude and longitude.

### Nest box monitoring and nestling condition

Each year, nest boxes were monitored prior to the arrival of Tree Swallows and then every two days throughout the breeding season (Appendix S1: Figure S1). This detailed monitoring allowed us to accurately estimate the dates of laying, incubation, hatching, and fledging, as well as record the number of eggs, nestlings, and fledglings of each breeding event. We caught and banded adult females during incubation and adult males during food provisioning. We were 99% and 80% successful, respectively, at capturing targeted adults. Once caught, adults received an aluminum US Geological Survey (USGS) band containing a unique identification code. At 12 days of age, nestlings also received a USGS band. To prevent premature fledging, following 16 days of age, the number of nestlings remaining within nest boxes were only counted, thus the exact date each nestling fledged is unknown.

### Insect samples and prey availability

Throughout the duration of this study, two insect traps were placed on each of the 40 farms (N=80). Traps were spaced at least 250 meters apart along the central portion of each nest box transect. The content of traps was collected every two days throughout each breeding season. Insect traps were a combined window/water-pan flight trap. Traps were made by bisecting two plexiglass sheets (11.5 cm x 30 cm), and then fixing them on top of a yellow bucket 15 cm deep and 21 cm in diameter (Rioux Paquette et al. 2013, Bellavance et al. 2018). Traps were then attached to a stake one meter above the ground and filled with approximately one liter of a saltwater and soap solution. The content of each trap was collected by straining out insects into a tube filled with 70% ethanol. Samples were then stored in closed boxes at room temperature until processing.

Our interest being in prey availability during the nestling rearing period, we focused the processing of insect samples collected between 1 June and 15 July, covering over 96% of breeding attempts (Appendix S1: Figure S1). The processing of insect samples started by first removing items and insects not found in the diet of Tree Swallows. This included removing gastropods, large spiders, June bugs (Coleoptera: Melolonthinae), large Lepidoptera, and bumblebees [(*Bombus* spp.), (Winkler et al. 2020)]. The remaining content of a trap was composed of a wide range of insects stemming from many different habitat types (Laplante 2013) and overlapped significantly with the insects found within food boluses provided to nestlings (Bellavance et al. 2018). In our system, 74% of the biomass and 77% of the insects found within food boluses provided to nestlings are Diptera (Bellavance et al. 2018 and this study), and this pattern is shared with other Tree Swallows study systems (McCarty and Winkler 1999a, Johnson and Lombardo 2000, Mengelkoch et al. 2004, Twining et al. 2018). We thus used the biomass of Diptera within each sample as a proxy of prey availability (using the number of Diptera resulted in qualitatively similar results; analyses not shown). We evaluated the performance of prey biomass and fitness models when treating prey availability as all insects or only Diptera (38% of the biomass from samples) and found models performed substantially worse when treating prey availability as the biomass of all insects (Appendix S2: Table S1). Moreover, we did not limit processing of Diptera to any specific size limit as nearly all major groups of Diptera from our system can be found within the food boluses of Tree Swallows (Bellavance et al. 2018). Once extracted, the number of individual Diptera was recorded and the sample was placed in an oven at 60°C for over 24 hours, ensuring no further change in biomass occurred. Once dried, samples were weighed without delay (± 0.0001 g). The dried biomass of Diptera (hereinafter referred to as Diptera biomass) from these samples thus represents an estimate of prey availability on a specific farm and year over the two days prior to the sample collection.

### Fitness and body condition proxies

We focused on four proxies of fitness and the body condition of fledglings; the proportion of a brood that fledged (fledging success), the duration of the nestling period, and the body mass and wing length of fledglings. Field methods did not allow us to define the fledging dates of specific nestlings. Consequently, the duration of the nestling phase was represented by the number of days between the maximum hatching date and mean Julian date between the first and last nestling to leave the nest. At 16 days of age, we measured nestling mass (± 0.01 g) and wing lengths (± 1 mm) using a wing ruler. Mean fledging age was 19.69 ± 1.77 (SD) and the mean number of days between the first and last nestling to leave the nest was 1.58 ± 1.08 (SD). We therefore assumed morphological measures at 16 days of age were representative of fledgling morphology (Zach and Mayoh 1982, McCarty 2001).

### Landscape composition

Landscape composition (i.e., the relative coverage of habitats composing a given landscape) focused on habitats Tree Swallows are hypothesized to use as indicators of nest-site selection, and the principal habitats composing landscapes throughout this system (Rendell and Robertson 1989, 1990, Winkler et al. 2020). The relative cover of habitats was calculated within 500 m of nest boxes. This spatial scale represents the area within which nestling-rearing female Tree Swallows spend ∼80% of their time (Elgin et al. 2020, Garrett et al. 2021) and is the largest extent for which exhaustive annual characterization of landscapes has been performed in our system since 2006 (Porlier et al. 2009). Moreover, Tree Swallows are often associated with aquatic habitats and will travel distances greater than 500 m to forage over them (Elgin et al. 2020, Garrett et al. 2021). However, the relative cover of aquatic habitats within 500 m of nest boxes and in these agricultural contexts was low 0.66% ± 1.07%. We therefore considered these habitats as being potentially influential to the fitness of Tree Swallows (Berzins et al. 2021) and calculated their relative cover at greater distances following the characterization of landscapes within 500 m. Parcels representing different habitats and agricultural fields within each 500-m buffer were delineated using orthophotos (scale 1:40 000) in QGIS (QGIS 2020). Content of parcels were then characterized *in situ* at the end of the breeding season but before harvest in order to facilitate their identification. This included characterizing which cultures, if any, were in agricultural fields. Habitats and cultures were then reclassified into one of five higher order categories. These higher order categories are forested, corn and soybean, forage fields (including hay fields, other grasses, alfalfa, clover, pastures, and old fields), and cereals (other than corn and soybean). We then calculated the mean percent cover of higher order habitats across the 10 nest boxes on each farm and for every year independently.

To obtain an integrative measure of the percent cover of all higher order habitats, defined as the landscape context, we used a robust principal components analysis (PCA) for compositional data (Filzmoser et al. 2009) to assign “site scores” to each of the farms during each year. Site scores were defined by the values along the components of the resulting compositional PCA of each farm during each year as fitted using the robCompositions package (Templ et al. 2011) in R version 3.6.2 (R Core Team 2019). This method takes into consideration the principle that values of compositional data are inherently related to one another given they must sum to one and resulted in the calculation of 440 different landscape contexts observed throughout the duration of the study (i.e., 40 farms x 11 years). Site scores were then assigned to each breeding attempt and insect sample and used in all subsequent analyses as they allow for comparative measurements of landscape contexts.

The first component (Comp.1) of the PCA explained 80.34% of the variance in landcover and correlated positively with corn and soybean and negatively with both forage fields and forest cover. The second component (Comp.2) explained 14.69% of the variance in landcover and correlated negatively with forage fields and positively with forest cover (Figure 2). The first two components of the PCA explained over 95% of the variance in our landscape characterization dataset. We attempted to keep models as simple as possible and because forest cover has been shown to be influential to the nest site selection and foraging behavior of this species (Rendell and Robertson 1989, Courtois et al. 2021, Garrett et al. 2021), we included only these two components to represent the landscape context. Landscapes expressed by maximizing Comp.1 and minimizing Comp.2 represent ones for which there is a mixture of corn, soybean, and cereals and denuded of forest cover, and are thus referred to as mixed-row crop landscapes. Landscapes expressed by minimizing Comp.1 and with negative Comp.2 values represent ones for which there is an abundance of forage and pasture fields with average forest cover and are thus referred to as forage landscapes.

**Figure 2:**
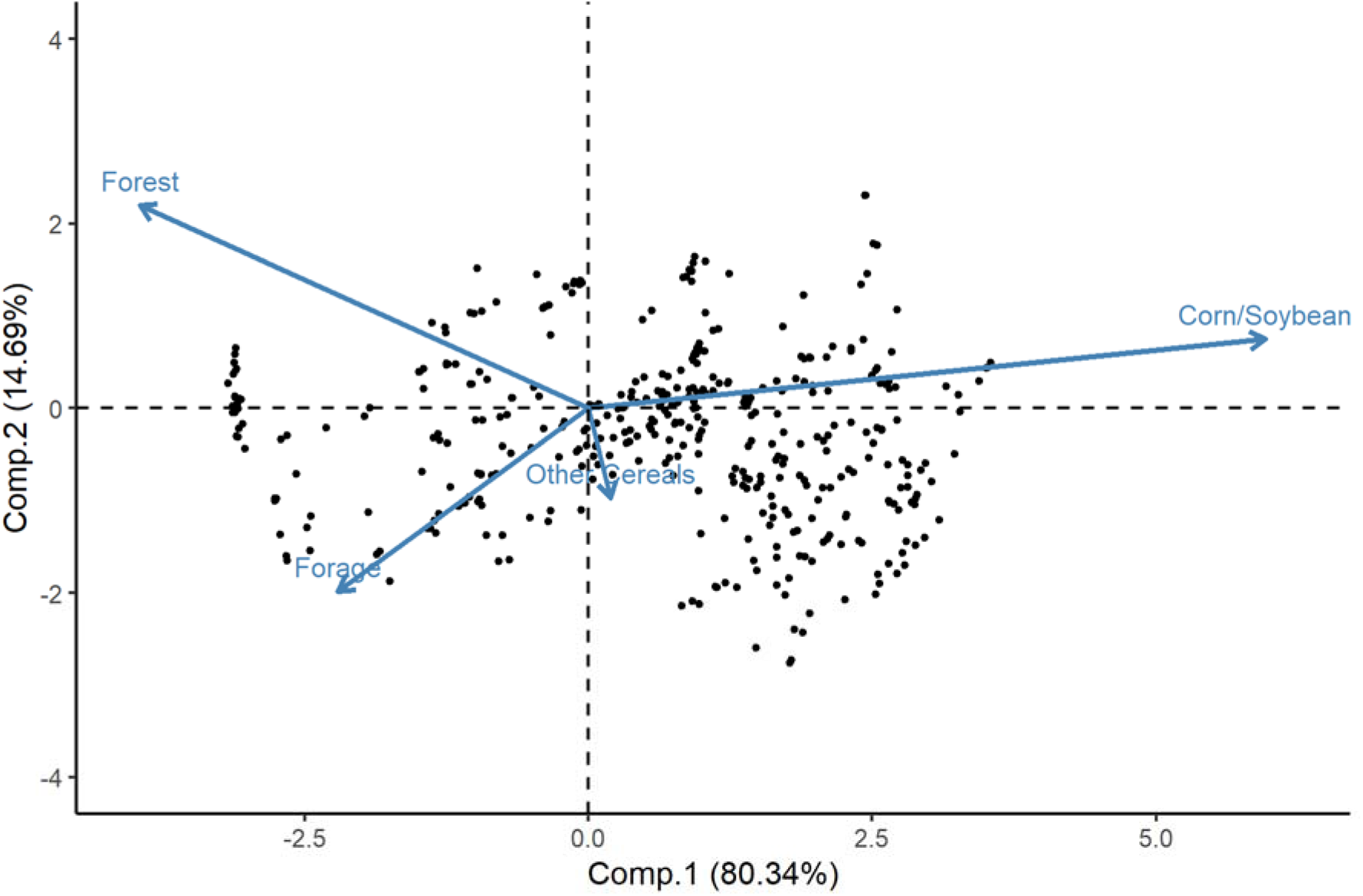
Robust compositional principal components analysis (PCA) of landscape habitat composition surrounding each of the 40 farms between 2006 and 2016. Arrows indicate the eigenvalue and loadings of each higher order habitat. Points within the background represent the site scores assigned to each farm and year combination and used to define landscape context (N=440 farms*years)

Finally, to calculate the relative cover of aquatic habitats, we performed a post-hoc model comparison determining the spatial extent to which the percent cover of open water was influential in predicting fledging success. Each model had an identical set of covariates and random effect structure (model: Base + Food * Land in Appendix S1: Table S1), varying only by the percent cover of open water calculated at increasing spatial extents between 500 m to 10 km, at 500-m increments. The model resulting in the lowest second-order Akaike information criterion (AICc; Burnham and Anderson 2002) occurred when considering the percent cover of open water within 3 km of a nest box. We thus included the percent cover of open water within 3 km around each breeding attempt as another landscape variable. Open water data were derived from the annual crop inventory produced by Agriculture and Agri-Food Canada (AAFC; AAFC 2018).

### Weather variables

Due to the ectothermic nature of insects, measurements of prey availability can be strongly influenced by weather conditions during each sampling event (Winkler et al. 2013, Shipley et al. 2020). Furthermore, Tree Swallow breeding success may also be influenced by weather conditions independently of its effects on prey availability through, for example, the induction of hypothermic conditions of nestlings (Ardia et al. 2010). We wanted to control for these effects by including parameters indicative of temperature and precipitation during insect sampling and during the pertinent time windows of each breeding attempt. We therefore collected hourly temperature with iButtons (model DS1922L; Embedded Data Systems, Lawrenceburg, Kentucky, USA) attached to the underside of a single nest box on each farm. Temperatures from data loggers placed on the outside of nest boxes were highly correlated to those recorded inside of nest boxes (*r* = 0.97), and therefore we used the former as they are likely more representative of the temperatures experienced by aerial insects and foraging adult Tree Swallows. Hourly temperatures were recorded throughout the entirety of each breeding season and started prior to the arrival of Tree Swallows onto the system. From these data, we derived daily summaries of the minimum, mean, and maximum daily temperatures. We measured total farm-specific precipitation every two days throughout the breeding season using a single pluviometer placed on each farm and recorded measurements to the nearest 0.5 ml during each farm visit.

### Statistical analyses

#### Seasonal influence of weather and landscape composition on prey availability

We assessed the influence of covariates on prey availability using generalized linear mixed-effects models (GLMMs). We assessed a series of models representing alternative hypotheses in order to evaluate if agricultural landcover is related to prey availability either additively or through interactions with seasonality and/or weather. We calculated the mean temperature values between the day of and day preceding the collection of insect samples, to ensure that weather covariates were representative of the estimate of prey availability. We treated precipitation the same way, as the bi-daily recordings already represented the accumulated precipitation during each insect sampling period. The conditional distribution of Diptera biomass was right-skewed, therefore GLMMs incorporated a Gamma distribution and a log link function. Fixed effects within competing models included the site scores (Comp.1 and Comp.2) and their interaction, Julian date, and weather variables associated with the collection of each sample. We treated the effect of Julian date as a second-degree polynomial, expecting insect abundance to reach a peak around the middle of the breeding season (Rioux Paquette et al. 2013, Bellavance et al. 2018). Ad-hoc model comparison using full models (model: Base + Rain * Land + Time * Land; Appendix S1: Table S1) highlighted that the most predictive temperature of Diptera biomass was the mean maximum temperature over the two-day period (AICc *w* = 0.94). We therefore treated temperature as the mean maximum during this period. Because we did not expect Diptera biomass to respond linearly with temperature, as it typically rises at an increasing rate with temperature before plateauing and subsequently declining, indicative of a third degree polynomial, we treated the effect of temperature as such (Winkler et al. 2013). Finally, preliminary exploration highlighted that the phenology of Diptera biomass varied markedly among both years and farms. We hence performed further ad-hoc model comparisons of two full models, differing only in the presence of a variable slope of Julian date, treated as a second-degree polynomial, across years and farms through random effects. AICc weights greatly favored this variable slope model (AICc *w* >0.99). All implementation of polynomial terms was with the poly() function in R.

Evaluating the strength of evidence for the hypothesis that prey availability is related to landscape context took an information theoretic and multimodel approach (Burnham and Anderson 2002). Alternative hypotheses were thus evaluated using AICc and the associated Akaike weight (*w*) of competing models. We hypothesized that the weather dependent nature of insects may result in the influence of the landscape context on prey availability being either masked by or vary with the timing of sampling and weather conditions around sampling time (i.e., in the two days prior to insect collection). We accordingly defined a set of six candidate models including a null model exempt of all fixed effects (Appendix S1: Table S1). Two models treated fixed effects as strictly additive, including one where prey availability varied only with sampling time and weather (Base), and one with the landscape context, while controlling for both sampling time and weather (Base + Land). We also considered models where the effect of landscape context on prey availability varied either with the sampling time (Base + Time * Land), the amount of precipitation (Base + Rain * Land) or both the sampling time as well as the amount of precipitation (Base + Rain * Land + Time * Land).

### Influence of prey availability and agricultural landcover on fitness and body condition proxies

We reduced analyses to first breeding attempts (90% of all attempts and ranging between 80% and 95% across years), as reproductive success can vary greatly between first and second attempts, and as second attempts occur nearly exclusively following the failure of a nest during laying or incubation. Fledging success was modeled as the number of fledglings over brood size (i.e., number of successes over trials) via GLMMs with a binomial distribution and logit link function (Douma and Weedon 2019). The duration of the nestling period as well as nestling body mass and wing length were modeled using linear mixed effects models (LMMs). We included year, farm and nest box identification as random factors for fledging success and duration of the nestling period. Since multiple observations of nestling body mass and wing length for each brood were observed and nesting brood identification within nest box identification led to convergence failures, we nested brood identification within farm identification, along with year, as random factors.

Model covariate summaries can be found in Table 1 and were averaged across a 12-day window post-hatching for fledging success and a 16-day window post-hatching for the duration of the nestling period, nestling body mass and wing length. Twelve days of age represents the period when Tree Swallow nestlings become homeothermic, reach peak body mass, and during which most nestling mortality occurs, thus representing the period where food availability is presumably most crucial (McCarty 1995, Lamoureux 2010, Houle et al. 2020). Estimates of prey availability during the time window of each breeding attempt was represented by predictions from generalized additive models (GAMs) in which raw values of prey availability (i.e., Diptera biomass) were regressed against the Julian date of sample collection for each farm and year separately. We modeled Diptera availability to capture the general phenology throughout each season and avoid biases caused by more punctual or local disturbances or phenomena such as the capture of an insect swarm. GAMs were fitted as a tensor product smoother using the mgcv package in R (Wood 2015) with an identical degree of smoothness (k=10) and using a Gamma distribution with a log link function. From these predictions, we calculated an estimate of the mean availability of prey during the respective temporal window of each fitness and body condition proxy for each breeding attempt.

**Table 1:** Mean (±SD) of covariates used to model response variables. Covariates for prey availability were averaged over two days prior to the collection of each insect sample, whereas those for fitness and body condition proxies were averaged over either a 12-day (fledging success) or 16-day (nestling period duration and nestling body mass and wing length) window post hatching (presented are values for fledging success). Dashes imply the covariates were not used in the modeling of the response variable. Model groups identify which covariates were present within models found in Appendix S1: Table S1.

| Covariate | Acronym | Unit | Prey availability | Fitness and body condition proxy | Model group |
| --- | --- | --- | --- | --- | --- |
| Julian day | JJ | (day) | 172.20 ± (12.21) | - | Time |
| Maximum temperature | MAX | (°C) | 25.61 ± (3.75) | 25.36 ± (2.31) | Base |
| Precipitation | RN | (ml) | 7.16 ± (11.13) | 7.44 ± (4.92) | Base |
| Site score component 1 | Comp.1 | - | 0.42 ± (1.77) | 0.18 ± (1.81) | Land |
| Site score component 2 | Comp.2 | - | -0.24 ± (0.91) | -0.18 ± (0.92) | Land |
| Open water | WT | (%) | - | 0.01 ± (0.01) | Base |
| Prey availability (Diptera dry biomass) | FDM | (g) | - | 0.03 ± (0.02) | Food |
| Hatching day | HD | (day) | - | 159.90 ± (5.76) | Base |
| Brood size | NN | (nestlings) | - | 4.83 ± (1.27) | Base |
| Brood days | BD | (summation of daily brood size) | - | 85.63 ± (18.19) | Base |

In order to evaluate our hypothesis regarding trophic links between agricultural landscapes and fitness and body condition proxies, we compared a set of six models, each representing biologically relevant alternative hypotheses predicting the outcome of each proxy and took an information theoretic and multimodel approach (Appendix S1: Table S1). Besides a null model only containing random effects, we started with a model including only confounding factors (Base). Confounding factors remained identical across all fitness and body condition proxies and were included to control for their known effects. Confounding factors included the mean maximum daily temperature (Winkler et al. 2013) and mean daily precipitation (Cox et al. 2019, 2020) during the same post hatching and proxy-specific temporal window as prey availability. We also included the morphological age of the breeding female (second year (SY) vs. after second year (ASY); Hussell 1983), brood size, and the percent cover of open water around a nest box. Finally, to account for seasonal variation in the species composition of Diptera (Bellavance et al. 2018), and the trend of seasonal declines in reproductive success observed by Tree Swallows (Winkler and Allen 1996), we also included a brood’s hatching date. In the case of nestling body mass and wing length, brood size was replaced with the summation of the daily brood size during the 16-day period (Brood days), as this accounts for intra-brood competition in the event of nestling mortality (Martin 1987). All model terms within Base were included in all subsequent models of the candidate set. We included three models hypothesizing the effect of prey availability and landscape context were influential in a strictly additive capacity, including a model with only prey availability (Base + Food), a model with only site scores and their interaction (Base + Land), and a model with both prey availability and site scores (Base + Food + Land). The final model is one hypothesizing that the effect of prey availability varied with landscape context and thus included an interaction between prey availability and both sites score values (Base + Food * Land).

In all analyses of both Diptera availability and fitness and body condition proxies, unless a top performing model could be determined (AICc *w* > 0.90), the effects of key individual variables, including interactions, were estimated via multi-model inference whereby model predictions were calculated by model-averaging with shrinkage and shown with their 95% unconditional confidence intervals (Burnham and Anderson 2002). All quantitative covariates were z-transformed, r-squares were calculated following Nakagawa and Schielzeth (2013), and all analyses were performed in R using the glmmTMB (Magnusson et al. 2020) and AICcmodavg (Mazerolle 2020) packages. Model validation included evaluation of normally distributed residuals (simulated), heteroskedasticity, and checks of variance inflation factors (VIF) following Zuur et al. (2009) and using the DHARMa package (Hartig 2020).

## Results

### Seasonal influence of weather and an agricultural gradient on prey availability

We collected and processed 15,916 insect samples, resulting in information on 8,614 farm visits over 11 years. Overall mean Diptera availability (± SD) was 0.030 ± 0.044 g per trap and per farm for each two-day sampling period. While it did not vary greatly between years (range of means: 0.019 g - 0.037 g), the variance within each year was high (range of SD: 0.023 g - 0.059 g; Appendix S1: Figure S2). Diptera availability peaked near Julian day 183 (June 24; Figure 3a), decreased following increasing precipitation, and rapidly increased with mean maximum temperature, peaking at 21.7°C (Table 2; Appendix S1: Figure S3). While the interaction between site score values indicated Diptera availability was highest within predominantly forested landscapes (range percent forest cover: 0.0% to 69.1%), it was marginally greater in predominantly forage landscapes than in ones dominated by corn and soybean fields (Figure 3f). Compared to forage landscapes, predicted Diptera availability was 14.0% lower in corn and soybean landscapes as well as mixed-row crop landscapes, and nearly 200% greater in forested landscapes. Furthermore, the interaction between site scores and Julian date highlighted that, at the start and end of the food provisioning period (Julian date 152 and 196, respectively), Diptera availability within corn and soybean landscapes was 59.2% and 62.9% lower than within forage landscapes (Appendix S1: Figure S4). While during the middle of the food-provisioning period (Julian date 174), predicted Diptera availability was only 5.2% lower within corn and soybean landscapes versus forage landscapes. Finally, we did not find strong support that the influence of precipitation on Diptera biomass varied with landscape context.

**Figure 3:**
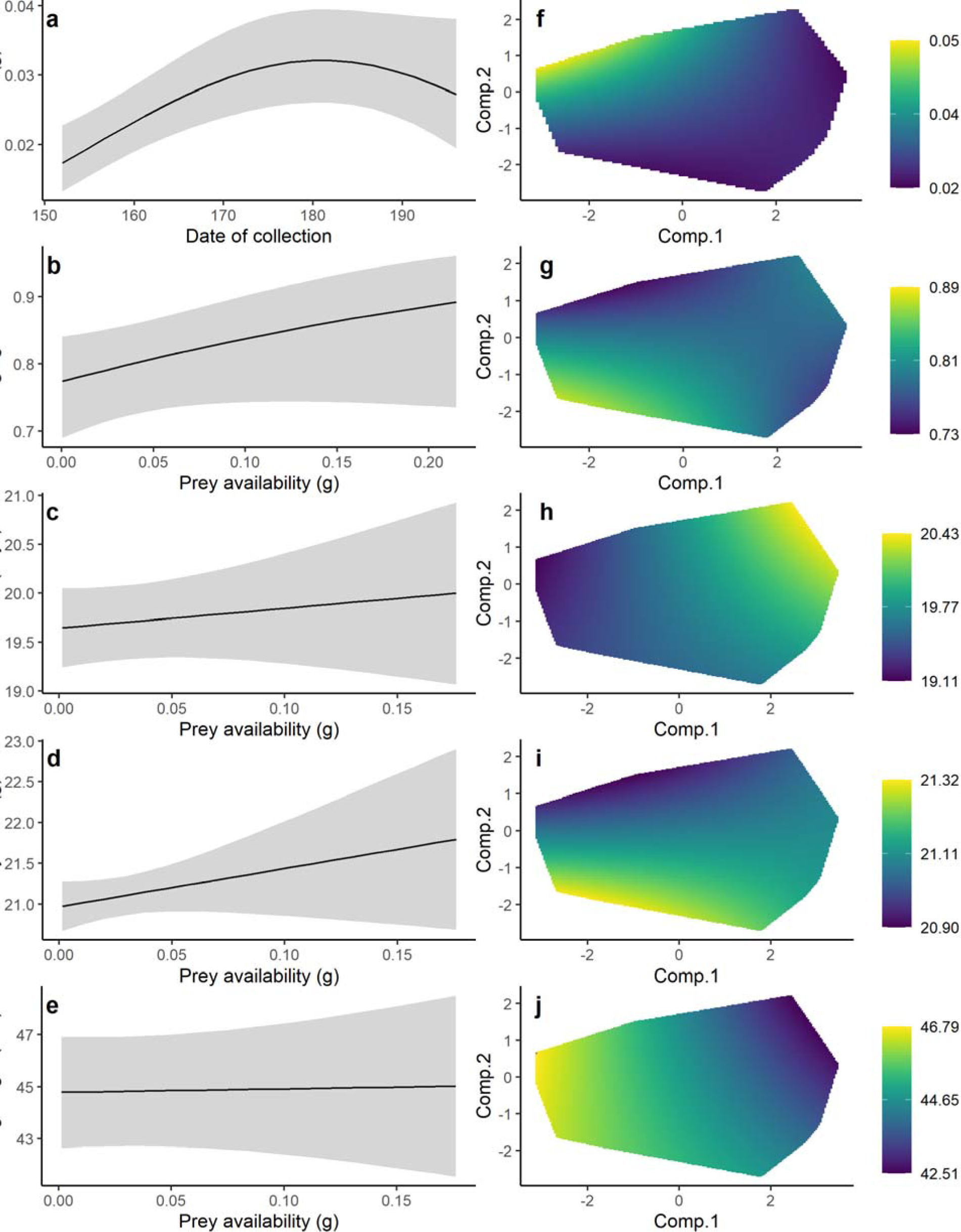
Predictions for the main covariates of interest. All predictions are unconditional predictions that have been model averaged, with shrinkage, according to Appendix S1: Table S1. Each row are predictions for the same response variable. Each plot within the first column, except for Figure a, is one fitness or body condition proxy plotted against the full range of observed prey availability. Figure a is the predicted Diptera biomass against the range of Julian dates. All error bars are unconditional 95% confidence intervals. Each plot within the second column is a prediction surface of each response variable against the first two components of the robust PCA describing landscape context. The prediction surface has been reduced to the convex hull of site scores observed within this study. The range of colors represent the minimum (deep blue) and maximum (yellow) predicted values of each response variable given site score values. Individual plots are the predicted Diptera biomass against Julian date of collection (a), the predicted fledging success (b), duration of the nestling period (c), body mass (d) and wing length (e) of nestlings at 16 days of age against the full range of prey availability. The predicted Diptera biomass (f), fledging success (g), duration of the nestling period (h), body mass (i) and wing length (j) of nestlings at 16 days of age against the first two components of the compositional PCA. Predictions and their covariates have been back transformed to either the response or unstandardized scale, respectively. All covariates not explicitly presented in figures were fixed to their mean values prior to predictions.

**Table 2.** Standardized coefficient estimates, and their 95% confidence intervals of the top model for each response variable. See Table 1 for definitions of covariates and Appendix S1: Table S1 for outcome of model selection. Coefficients in bold indicate confidence intervals not overlapping zero. Numbers next to acronym refer to the order of the polynomial term.

| Covariate | Biomass | Fledging success | Duration | Body mass | Wing length |
| --- | --- | --- | --- | --- | --- |
| FDM | - | **0.11**<br>(0.02, 0.19) | - | **0.10**<br>(0.01, 0.20) | - |
| Comp.1 | **-0.11**<br>(-0.17, -0.06) | -0.02<br>(-0.10, 0.06) | **0.17**<br>(0.08, 0.26) | - | **-0.58**<br>(-0.92, -0.24) |
| Comp.2 | **0.11**<br>(0.04, 0.17) | **-0.15**<br>(-0.28, -0.01) | 0.05<br>(-0.11, 0.21) | - | -0.20<br>(-0.81, 0.40) |
| Comp.1:Comp.2 | **-0.04**<br>(-0.07, -0.01) | **0.08**<br>(0.01, 0.15) | 0.04<br>(-0.05, 0.12) | - | -0.08<br>(-0.41, 0.25) |
| Comp.1:FDM | - | **-0.05**<br>(-0.10, -0.01) | - | - | - |
| Comp.2:FDM | - | **0.08**<br>(0.00, 0.17) | - | - | - |
| WT | - | **0.24**<br>(0.09, 0.39) | -0.08<br>(-0.24, 0.07) | 0.08<br>(-0.08, 0.23) | 0.49<br>(-0.10, 1.07) |
| HD | - | **-0.28**<br>(-0.35, -0.20) | -0.05<br>(-0.15, 0.06) | **-0.26**<br>(-0.36, -0.16) | 0.11<br>(-0.27, 0.49) |
| MAX | - | **0.30**<br>(0.21, 0.39) | **-0.33**<br>(-0.45, -0.21) | 0.04<br>(-0.08, 0.17) | **1.82**<br>(1.35, 2.30) |
| RN | **-0.07**<br>(-0.08, -0.05) | **-0.11**<br>(-0.20, -0.03) | 0.03<br>(-0.08, 0.15) | -0.12<br>(-0.23, 0.00) | 0.08<br>(-0.38, 0.54) |
| AGSY | - | **-0.25**<br>(-0.44, -0.06) | **0.36**<br>(0.09, 0.63) | 0.01<br>(-0.25, 0.28) | **-1.38**<br>(-2.39, -0.38) |
| NN | - | **-0.18**<br>(-0.25, -0.11) | **0.26**<br>(0.18, 0.34) | - | - |
| BD | - | - | - | **-0.22**<br>(-0.29, -0.14) | -0.06<br>(-0.35, 0.22) |
| JJ1 | **17.77**<br>(6.40, 29.14) | - | - | - | - |
| JJ2 | **-10.69**<br>(-20.31, -1.06) | - | - | - | - |
| JJ1:Comp.1 | 0.40<br>(-2.99, 3.79) | - | - | - | - |
| JJ2:Comp.1 | **-5.44**<br>(-8.19, -2.68) | - | - | - | - |
| MAX1 | **-11.86**<br>(-14.33, -9.39) | - | - | - | - |
| MAX2 | **-9.72**<br>(-11.92, -7.52) | - | - | - | - |
| MAX3 | 1.26<br>(-0.79, 3.31) | - | - | - | - |

### Fledging success

In total, we monitored the breeding activity of 1,897 breeding attempts producing at least one nestling across the 40 farms and 11 breeding seasons. Overall mean fledging success (± SD) was 0.74 ± 0.38 per brood and ranged between 0.63 and 0.88 among years, with the exception of the 2016 breeding season, where mean fledging success was 0.49 due to a period of prolonged cold temperatures in June resulting in many brood failures (Appendix S1: Figure S2). As expected, fledging success increased with prey availability (Table 2, Figure 3b). An increase of 17.0% in fledging success was predicted between broods experiencing the minimum and maximum observed levels of prey availability. A residual effect of land cover indicated that, under average prey availability, fledging success was greatest within landscapes minimizing site score values, and thus lowest within more agro-intensive landscapes (Figure 3g). Compared to forage landscapes, fledging success was 10.3% lower in corn and soybean landscapes and 13.3% lower within mixed-row crop landscapes. Interaction terms between site scores and prey availability indicated under low prey conditions (half of the mean), fledging success within corn and soybean landscapes or mixed-row crop landscapes was 8.6% and 9.7% lower than in forage landscapes, respectively. Furthermore, under typically high levels of prey availability (twice the mean), fledging success within these same two agro-intensive landscapes was 13.4% and 20% lower than in forage landscapes, respectively (Appendix S1: Figure S5). Hatching date, younger aged females (SY), brood size, and mean precipitation were all negatively related to fledging success (Table 2). Both the mean maximum daily temperature and percent cover of open water were positively related to fledging success (Table 2).

### Nestling duration

We measured the duration of the nestling period for 1,556 broods from which at least one nestling fledged. Mean nestling duration (± SD) was 19.7 ± 1.8 days per brood and ranged between 18.9 days and 21.1 days across years (Appendix S1: Figure S2). While we found no effect of prey availability on nestling duration, the latter was 6.9% longer between maximum and minimum Comp.1 scores and decreased slightly along Comp.2 (Figure 3h). Nestling duration was thus longest within landscapes dominated by row crops and shortest within those occupied by a mixture of forage fields and tree cover. We found that nestling duration increased with both the younger age of the female (SY) and brood size but decreased with higher mean maximum daily temperature. However, we found no strong relationship with hatching day, mean precipitation, or percent cover of open water (Table 2).

### Nestling body mass

We weighed 6,011 nestlings from 1,400 broods at 16 days of age that survived to fledging. Mean nestling body mass (± SD) was 21.2 ± 2.0 g, ranging between 20.4 g and 21.5 g across years (Appendix S1: Figure S2). We found that nestling body mass increased with prey availability (Figure 3d). An increase of 3.9% in nestling body mass was predicted between the minimum and maximum observed levels of prey availability. After controlling for prey availability, we observed no residual effect of landscape context on body mass (Figure 3i). We further found nestling body mass decreased with high levels of precipitation, throughout the breeding season and with increasing number of brood days. However, nestling body mass was not related to the percent cover of open water or age of breeding females (Table 2).

### Nestling wing length

We measured the wing length of 6,065 nestlings from 1,413 broods at 16 days of age that survived to fledging. Mean nestling wing length (± SD) was 45.8 ± 7.0 mm, ranging between 38.7 mm and 49.2 mm across years (Appendix S1: Figure S2). We found wing length was unrelated to prey availability (Figure 3e). However, we observed a residual effect of landscape context on wing length whereby wing length was up to 10.1% longer within forage landscapes compared to corn and soybean landscapes (Figure 3j). We further found wing length increased with increasing mean maximum daily temperatures and was shorter for nestlings of young (SY) mothers and with increasing brood days. However, no relationship was observed between wing length and either precipitation, percent cover of open water, or hatching date (Table 2).

## Discussion

Peak prey availability for Tree Swallows occurred in landscapes dominated by forest cover and a mixture of agricultural habitats, suggesting greater landscape heterogeneity results in greater prey availability for this species throughout southern Québec’s gradient of agricultural intensification. We also observed that prey availability is an important determinant of Tree Swallow reproductive success in this geographical context. Indeed, greater prey availability was associated with higher fledging success and greater nestling body mass but unrelated to the time taken by nestlings to fledge or their wing length upon fledging. Interestingly, we found landscape context was associated with fledging success, nestling duration, and nestling wing length only after controlling for the effect of prey availability. This residual effect of landcover suggests other factors related to the agricultural landscape context affect breeding Tree Swallows.

### Seasonal influence of weather and agricultural landcover on prey availability

The availability of prey was marginally greater in forage dominated landscapes than within corn or soybean and mixed-row crop landscapes. These results are in agreement with those found in the literature and our own predictions suggesting aerial insect abundance would be greater within less agro-intensive habitats (Wickramasinghe et al. 2004, Evans et al. 2007, Grüebler et al. 2008). Our finding that prey availability was greatest within agricultural areas harboring elevated forest cover is likely explained by habitat heterogeneity resulting in a more abundant and diverse community of Diptera (Benton et al. 2003, Fahrig et al. 2011). Perhaps the most interesting finding was that the phenology of prey availability as Diptera varied across this agricultural gradient. Diptera biomass in the middle of the breeding season was similar between forage dominated landscapes and landscapes dominated by more agro-intensive cultures. However, at the beginning and ends of the breeding season prey availability was greater in less agro-intense areas. This pattern was observed in other studies from this system yet on much shorter sequences of years (Rioux Paquette et al. 2013, Bellavance et al. 2018). A possible cause of this spatio-temporal variation in Diptera biomass is that the phenology and species composition of Diptera varied across the agricultural gradient. Such a variation in species composition was indeed observed in previous work from this system (Bellavance et al. 2018 and Appendix S2: Figure S1-5). Tree Swallows, similar to many other passerine species, display seasonal declines in reproductive success thought to be related to changes in the availability or adequacy of resources (Winkler and Allen 1996, Nooker et al. 2005, Bourret et al. 2015). Therefore, the landscape driven differences in the phenology of food resources reported here suggests that the severity of seasonal declines in reproductive success of farmland birds could be landscape dependent.

### Effects of prey availability on fitness and body condition proxies

Fledging success and nestling body mass were positively related to prey availability. These findings support previous studies documenting similar relationships stemming from artificial changes in local food availability (Boutin 1990, Granbom and Smith 2006, Robb et al. 2008, Ruffino et al. 2014). For instance, multiple lines of evidence suggest declines in fitness proxies and changes in foraging behaviors of avian species following the decrease of prey availability through the application of various agro-chemical agents (Martin et al. 1998, 2000, Brickle et al. 2000, Hart et al. 2006, Poulin et al. 2010). Our results also agree with previous evidence from other swallow populations observing unaltered fluctuations in prey availability being positively related to fledging success and nestling condition (Quinney et al. 1986, McCarty and Winkler 1999b, Winkler et al. 2013, Teglhøj 2017, Twining et al. 2018). Yet, our results run counter to recent studies from other swallow populations where measurable variation in aerial insect prey was suggested to have no impact on seasonal reproductive success and morphological condition of offspring (Dunn et al. 2011, Imlay et al. 2017, McClenaghan et al. 2019). A frequent claim of these studies is that prey availability was overly abundant during periods of observation, and thus other resources limited fitness, highlighting the importance of conducting food availability studies across a broad enough range of habitat qualities to reduce the availability of resources below such a critical threshold (Fazey et al. 2005). We found evidence that, within agricultural systems, the growth of Tree Swallow populations could be food limited. However, to address the role of prey availability on the population dynamics of this species more fully, the availability of food resources will also need to be quantified with reference to both clutch size and egg condition, as these factors partly determine brood size and fledgling number (Meijer et al. 1989, Ardia et al. 2006, Gow et al. 2019). Indeed, food resources may limit clutch size through a reduction in resource allocation (Wilson et al. 2017) and impact incubation capacity and hatching by regulating the duration and time between foraging bouts (McCarty 1995, Monaghan and Nager 1997, Thomson et al. 2007).

We found no indication that wing length was related to the availability of prey, perhaps suggesting that prey availability during this study was adequate for allocating resources towards the growth of flight feathers. We also found that the duration of the nestling period was unrelated to prey availability, a result running counter to that of other swallow studies (Teglhøj 2017). One plausible explanation is that the supply of food resources was sufficient to meet the basal metabolic demand for the rate of nestling growth given the observed brood sizes (Leonard 2000, Stodola et al. 2010). The duration of the nestling period is likely a function of the nestlings’ growth rate given that nestlings should remain within nests until reaching a sufficient condition (Nilsson and Svensson 1993, Seki and Takano 1998, Rossmanith et al. 2007). Recognizing this, the lack of a relationship between the duration of the nestling period and prey availability leads us to suggest that, in our system, the duration of the nestling period may be driven by something other than the accumulation of food resources.

### Residual effects of agricultural landcover on fitness and body condition proxies

An important finding of our study is that relationships between fitness and body condition proxies and agricultural landcover was observed even after controlling for prey availability. Fledging success was lower, nestling period durations were longer, and nestling wing lengths were indeed shorter within areas dominated by large-scale row-crop monocultures, principally corn and soybean. Among the several hypotheses explaining such results, we discuss three non-mutually exclusive alternatives whereby differences in landscape context could lead to variation in the contamination of food by pesticides, in the nutritive value of food, and in the rate at which food is provisioned.

During the years covered by this study, areas intensively-managed for growing corn and soybean harbored elevated levels of pesticides in their surface waters, notably several herbicides (e.g., glyphosate, atrazine, S-metolachlor, imazethapyr) and neonicotinoid insecticides [e.g.,. clothianidin, thiacloprid, thiametoxam, (Giroux 2019, Montiel-León et al. 2019)]. Many of those pesticides were also found, even simultaneously, in the insects fed to nestlings by Tree Swallows in our system (Poisson et al. 2021) and may hence have caused lethal and sublethal toxicological effects on nestlings. For instance, many pesticides, such as the neonicotinoids, can alter the neurological, endocrine and immunological systems of non-target animals, including vertebrates, via skin contact, breathing or food/water consumption (Grue et al. 1997, Mayne et al. 2005, Mineau and Palmer 2013, Gibbons et al. 2014, Lopez-Antia et al. 2015). Such disruptions can induce, among many observed effects, anorexia and reduced mass gain, and even result in lower nestling survival, delayed fledging, and lower fledging mass, irrespective of the amount of food provided (Grue et al. 1997, Eng et al. 2017, 2019, Addy-Orduna et al. 2019). Pesticide exposure of Tree Swallow broods within areas dominated by corn and soybean may thus have been severe enough to result in nestling mortality or retarded nestling growth. Moreover, this may explain the lack of relationship between prey availability and the duration of the nestling period we observed, because exposure to some pesticides can also lessen parental care in birds (Grue et al. 1997, Custer 2011, Mineau and Palmer 2013, Gibbons et al. 2014).

Variation in landscape contexts may also have resulted in Tree Swallows feeding on diets differing not only in prey species composition, but also in nutritive value. Indeed, the diet of Tree Swallows varies across the agricultural intensification gradient covered by our study area (Bellavance et al. 2018; see also Nocera et al. 2012, Michelson et al. 2018). In the period during which Tree Swallows provide food to nestlings, they consume a diet with a greater biomass of insects with aquatic larval stages (e.g., Ephemeroptera and Diptera: Nematocera) in landscapes devoted mostly to forage crops and pastures (Bellavance et al. 2018 and Appendix S2: Figure S1-5). This is despite these insects representing a minute portion of what is available across the agricultural gradient studied here (Bellavance et al. 2018 and Appendix S2: Figure S1-5). Increased provisioning of these types of insects may lead to a more nutritive diet in less intensively cultivated landscapes as they typically contain high levels of highly unsaturated omega-3 fatty acids (HUFA), which have been found to improve the mass gain, immunocompetence, and fledging success of Tree Swallow nestlings (Twining et al. 2018). That said, insects with an aquatic larval stage can also be loaded with various environmental contaminants, such as pesticides and heavy metals, that may impair the breeding behavior and success of swallows as well as their survival (Custer 2011, Alberts et al. 2013, Twining et al. 2021). The costs and benefits of feeding on aquatic insects rich in HUFA within an agricultural context will thus require further attention before we can determine its real impact on Tree Swallow fitness components and demography.

Landscapes dominated by row crop monocultures express landscape simplification in which large swaths of areas are occupied by only a handful of habitats (Benton et al. 2003). This phenomenon may make finding and exploiting suitable food resources more difficult for animals relying upon “residual” marginal habitats as landscape simplification lowers the functional connectivity of agricultural landscapes (Hinsley 2000, Bélisle 2005, Rainho and Palmeirim 2011). If the amount and quality of suitable foraging habitats are reduced and disconnected, we may expect a negative impact of such landscape simplification on nestling growth if parents cannot compensate by increasing foraging effort. There is currently insufficient evidence to evaluate if differing compositions of agricultural landscapes consistently impact foraging adults negatively, yet the evidence available does suggest this is the case (Poulin et al. 2010, Staggenborg et al. 2017), albeit weakly in Tree Swallows (Stanton et al. 2016). In our study system, we observed that the biomass of food boluses delivered to nestlings does not vary with landscape context, yet their delivery rate is lowest within landscapes dominated by row crops (Garrett et al. 2021). Assuming parents are maximizing provisioning rate of food items to their offspring, given the availability of prey, this may provide evidence that adults are having to spend more time foraging within these types of landscapes (Charnov 1976, Pyke et al. 1977). If sustained, this increased foraging effort could come at a cost to parents and lead to carry-over effects eventually impacting their future fitness (Saino et al. 1999, 2018, Harrison et al. 2011, Stanton et al. 2017).

## Conclusion

Taken together, our results provide evidence that food availability during the nestling period has a positive effect on fitness components of Tree Swallows breeding in farmlands. Yet, more importantly, after controlling for food resources, a residual effect of agricultural landcover persisted. The interactive effect of prey availability and agricultural intensification on fitness and body condition proxies likely means that, for breeding attempts within less agro-intensive areas, increased prey availability has a positive influence. Yet, similar increases in prey availability within more intensively cultivated landscapes potentially result in an increased toxicological load of food to nestlings as well as a reduced dietary quality. Future work should expand on the relationship between nestling condition, fledging success, and post-fledgling survival in landscapes harboring elevated levels of agricultural habitats. More specifically, it is unknown how the availability of food resources early in the breeding season is influenced by agricultural land cover and if this relationship impacts settlement patterns and clutch size (Rendell and Robertson 1990, Orians and Wittenberger 1991, Grüebler et al. 2010, Robillard et al. 2013). However, it is certain that these are important features of the seasonal reproductive success and the population dynamics of aerial insectivores. Furthermore, nestlings reared under reduced prey availability conditions may be unfit for post-fledging survival or pre-migration fueling (Saino et al. 2012, Jones et al. 2017). For example, the body condition of nestling Barn Swallows (*Hirundo rustica*) is positively correlated to post fledging survival and fledglings in better condition initiated migration sooner (Evans et al. 2020). The poor body condition expressed by fledglings within more agro-intensive areas may thus result in elevated post-fledging mortality, a field readily understudied (Cox et al. 2014) yet a key part in the annual cycle of this species and another missing link in the understanding of aerial insectivore declines.

## Supporting information

AppendixS1

AppendixS2

## Acknowledgments

We are indebted to the farm owners who kindly accepted to partake in our long-term study since 2004. Sincere thanks to the many graduate students, field and lab assistants who helped collect, enter, and proof data throughout the years, notably with respect to insect sample processing. This work was conducted under the approval of the animal care committee of the Université de Sherbrooke and was financially supported by Natural Sciences and Engineering Research Council of Canada (NSERC) discovery grants to FP, DG and MB and a Steacie fellowship to FP, two team research grants from the Fonds de recherche du Québec—Nature et technologies (FRQNT) to FP, DG and MB, by the Canada Research Chairs program to FP and MB, as well as by New Opportunities Funds of the Canadian Foundation for Innovation (FCI) to FP, DG and MB, the Canadian Wildlife Service of Environment and Climate Change Canada, and the Université de Sherbrooke.

