## AppendixS1 for "Combined influence of food availability and agricultural intensification on a declining aerial insectivore"

Appendix S1: Supplemental tables and figures

Table S1: Results of the model selection process, including details regarding the model groups found within each candidate model. Composition of covariates within each model group are found in Table 1. Rain * Land and Time * Land includes interaction terms with only Comp.1. Food * Land includes the two-way interaction terms between FDM and both Site scores. Modeling was via GLMMs with a Gamma distribution and log link function for Diptera biomass (N=15,916), GLMMs with a binomial error distribution and a log link function for fledging success (N=1,897), and LMMs for the duration of the nestling rearing period (N=1,556), as well as both the body mass (N=6,011) and wing length at 16 days of age (N=6,065). All models received year and farm as random effects. Nest box ID was also included in both the fledging success and the duration of the nestling period as a random effect. Brood ID was also included in both the body mass and wing length as a random effect.

| Response | Candidate model | K | *w* | R2m |
| --- | --- | --- | --- | --- |
| Biomass | Base + Time * Land | 25 | 0.73 | 0.06 |
|  | Base + Rain * Land + Time * Land | 26 | 0.27 | 0.06 |
|  | Base + Land | 23 | 0.00 | 0.06 |
|  | Base + Rain * Land | 24 | 0.00 | 0.06 |
|  | Base | 20 | 0.00 | 0.03 |
|  | Null | 14 | 0.00 | 0.00 |
| Proportion | Base + Food * Land | 16 | 0.72 | 0.05 |
|  | Base + Food + Land | 14 | 0.12 | 0.05 |
|  | Base + Land | 13 | 0.07 | 0.05 |
|  | Base + Food | 11 | 0.05 | 0.05 |
|  | Base | 10 | 0.04 | 0.04 |
|  | Null | 4 | 0.00 | 0.00 |
| Duration | Base + Land | 14 | 0.46 | 0.08 |
|  | Base + Food + Land | 15 | 0.35 | 0.08 |
|  | Base + Food * Land | 17 | 0.18 | 0.08 |
|  | Base + Food | 12 | 0.01 | 0.06 |
|  | Base | 11 | 0.00 | 0.06 |
|  | Null | 5 | 0.00 | 0.00 |
| Body mass | Base + Food | 12 | 0.40 | 0.03 |
|  | Base + Food + Land | 15 | 0.28 | 0.04 |
|  | Base | 11 | 0.13 | 0.03 |
|  | Base + Food * Land | 17 | 0.10 | 0.04 |
|  | Base + Land | 14 | 0.10 | 0.03 |
|  | Null | 5 | 0.00 | 0.00 |
| Wing length | Base + Land | 14 | 0.45 | 0.06 |
|  | Base + Food * Land | 17 | 0.36 | 0.07 |
|  | Base + Food + Land | 15 | 0.17 | 0.06 |
|  | Base | 11 | 0.02 | 0.05 |
|  | Base + Food | 12 | 0.01 | 0.05 |
|  | Null | 5 | 0.00 | 0.00 |

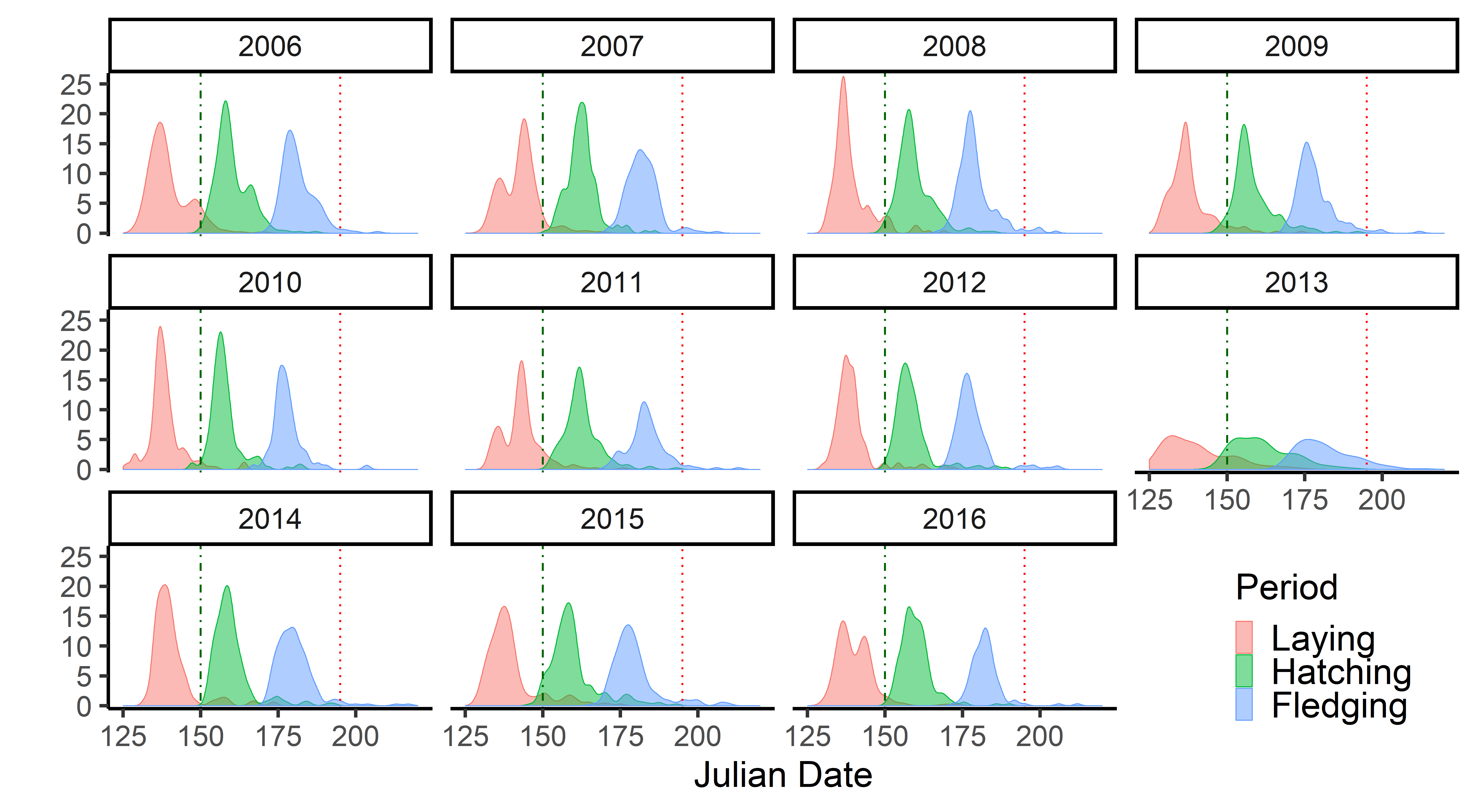

Figure S1: Histogram of the primary breeding periods experienced during each breeding season for Tree Swallow nesting in southern Québec between 2006 and 2016. Julian date is the number of days past 1 January. Vertical green (dashed) and red bars (dotted) represent the start and finish of the insect processing period, respectively.

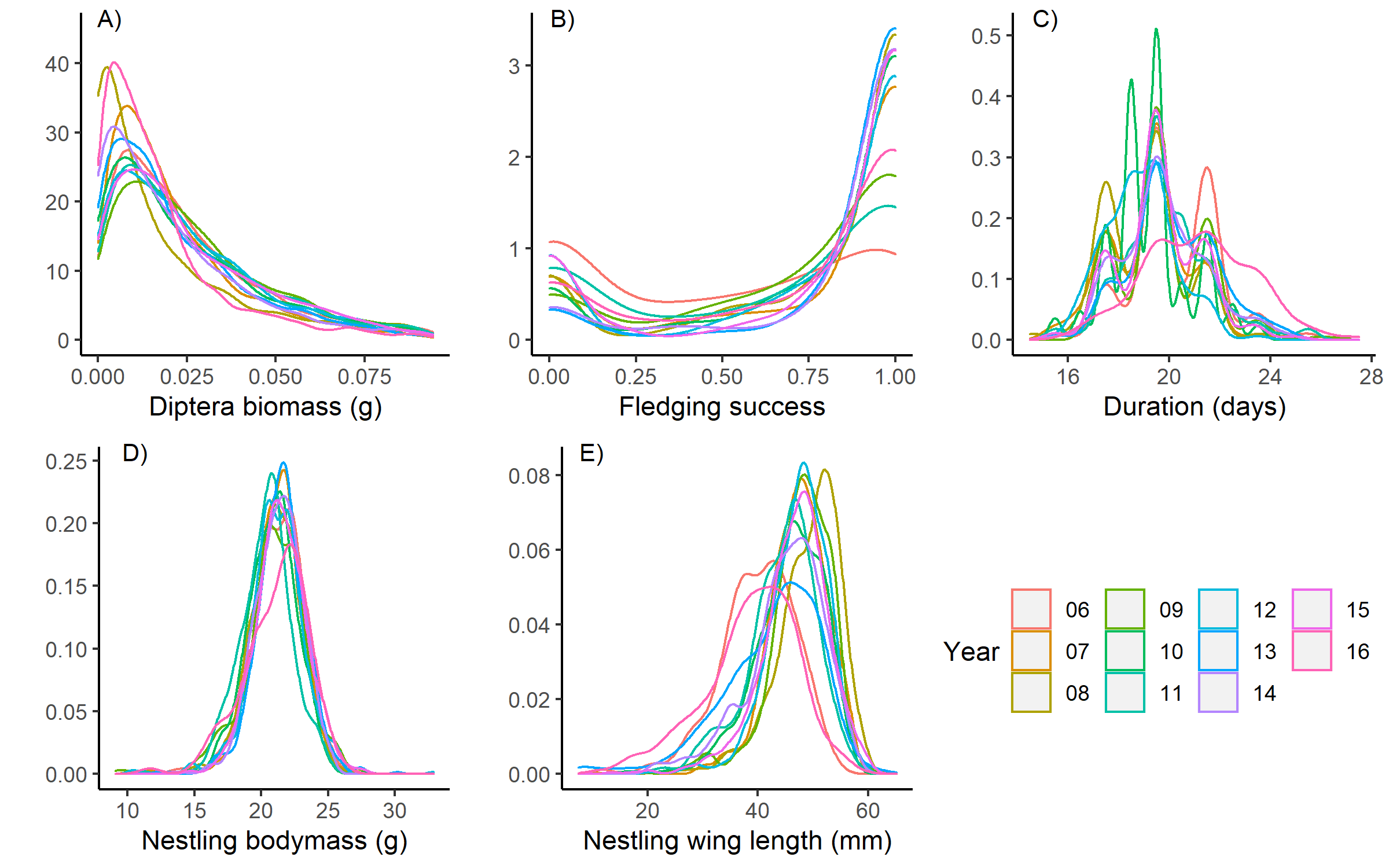

Figure S2: Kernel density estimate plot of each response variable throughout each year (2006-2016) of a study on the breeding ecology of a Tree swallow population distributed along a gradient of agricultural intensification in southern Québec, Canada. A) Diptera biomass (reduced to the lower 95% of values, (± 0.0001 g)), dried biomass of Diptera from two insect traps on each farm and collected every two days between 1 June to 15 July (representing over 95% of the first breeding attempts), B) Fledging success, the proportion of a brood that fledged (i.e. the number of fledglings/brood size), C) The duration of the nestling period, the difference between the mean Julian dates of the brood fledging and hatching, D) Nestling body mass (± 0.01 g), measured at 16 days of age, E) Nestling wing length (± 1 mm), measured at 16 days of age.

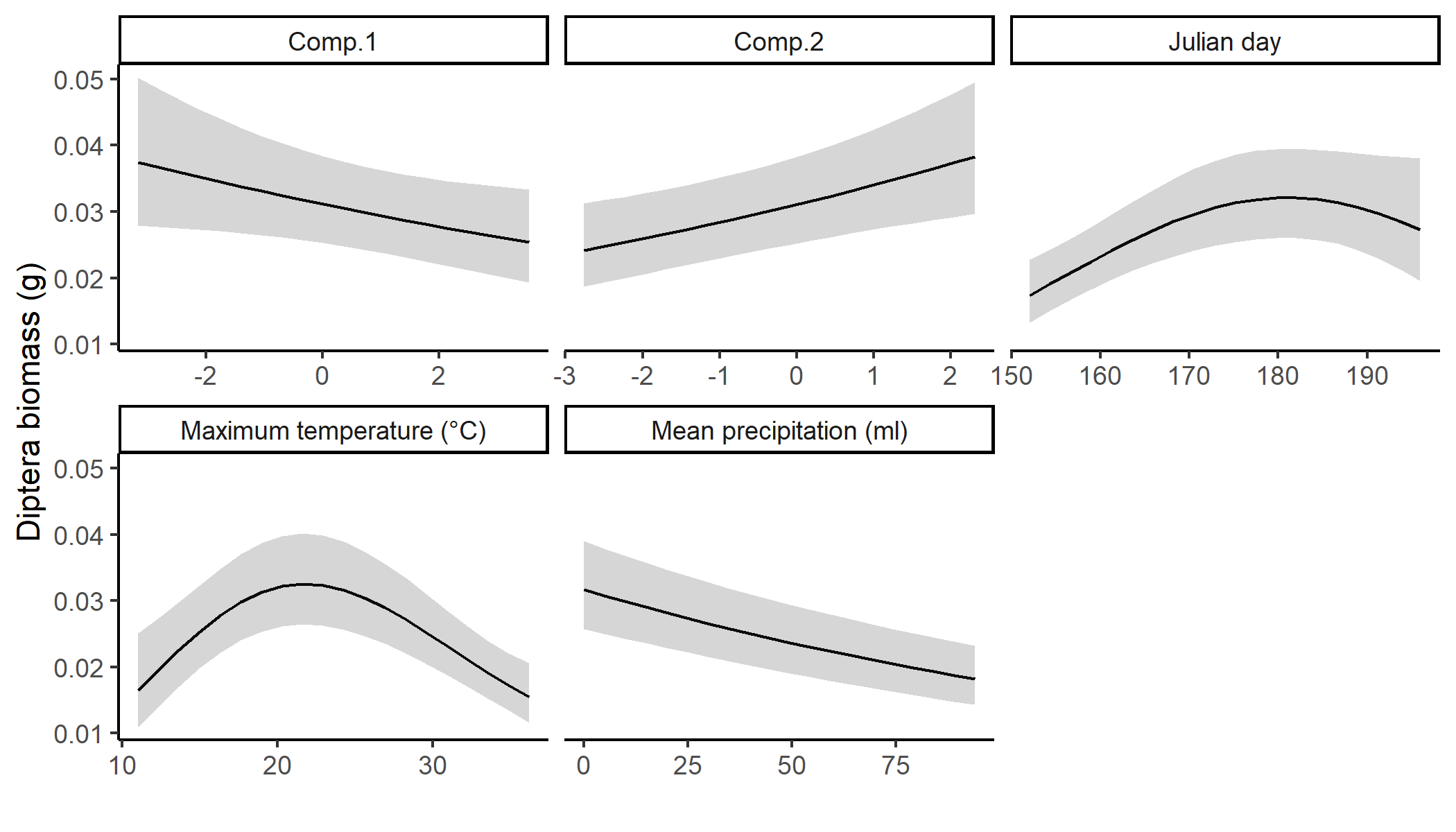

Figure S3: Predicted prey availability (Diptera dry biomass) with unconditional 95% confidence intervals against each covariate of interest. In each case, the values of all other covariates have been set to their mean and the predictor under consideration has been back transformed onto its original scale.

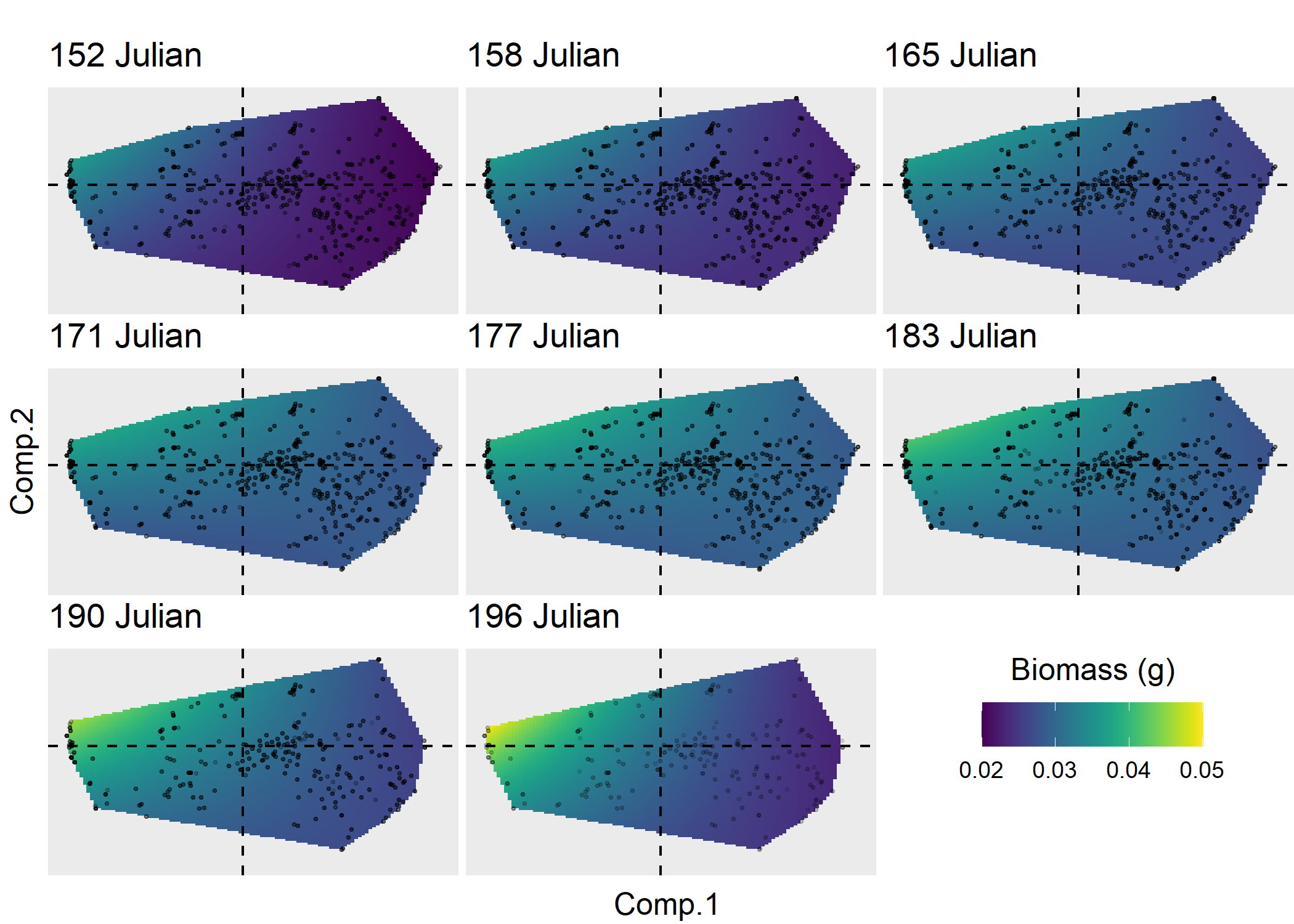

Figure S4: Predicted prey availability (Diptera dry biomass) against the first two components of the robust PCA describing landscape context for increasing levels of Julian date. Predictions and their covariates have been back transformed to either the response or unstandardized scale, respectively. Vertical and horizontal hashed lines represent the zero values for both components. Each prediction surface represents a different Julian date group. Julian date groups are the samples collected during different Julian dates between 152 to 196 for 6-day increments. Thus, raw values (points within background) represent the sites scores for the distribution of samples within each Julian date group. Prediction surfaces of each Julian date group were produced by first calculating the minimum Julian date, delineating out a bounding box of extreme site scores, and then deriving predictions based on these values, while keeping all other covariates at their mean. The range of colors represent the minimum (deep blue) and maximum (yellow) predicted values of each response variable given site score values.

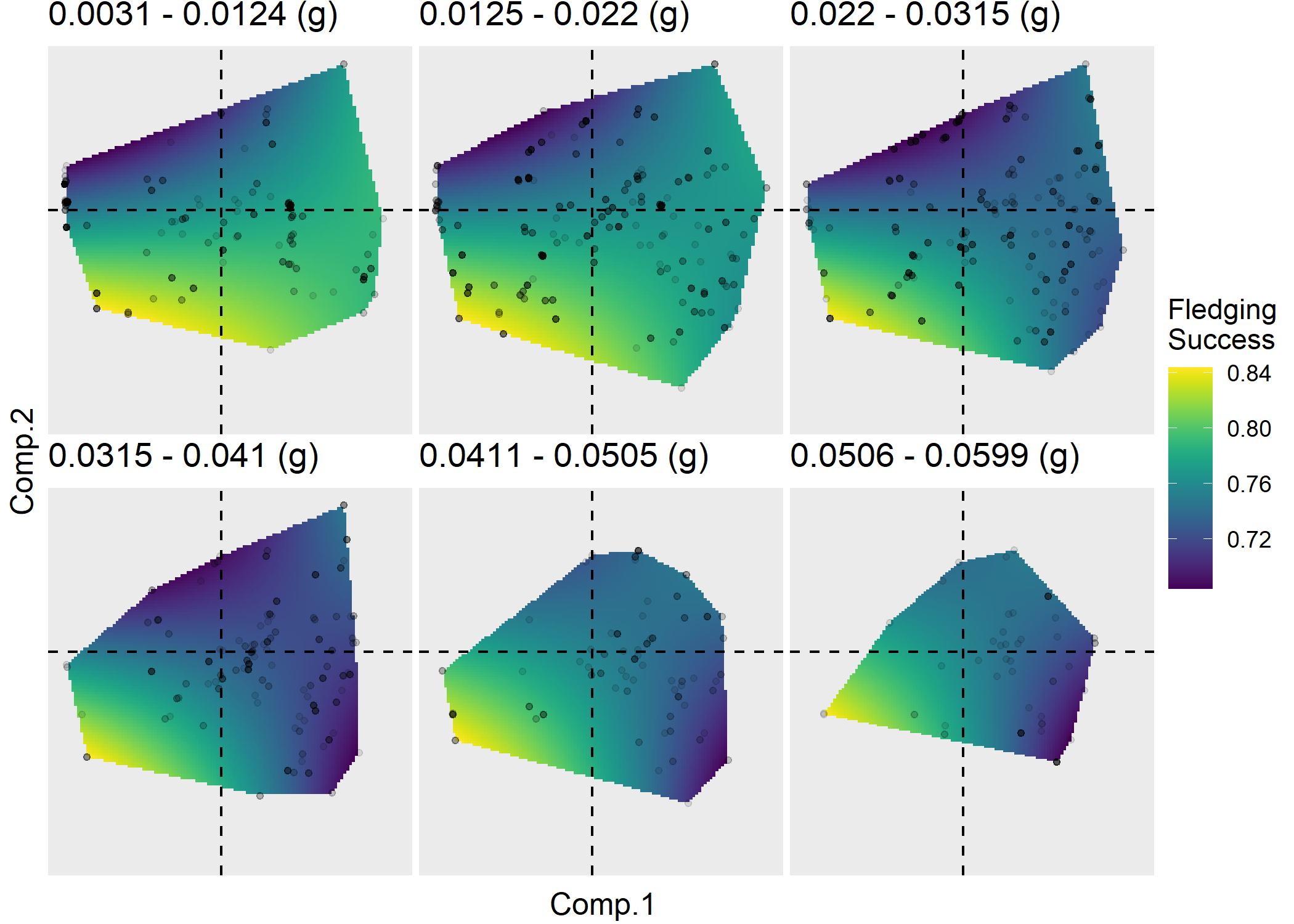
 Figure S5: Predicted fledging success against the first two components of the robust PCA describing landscape context for increasing levels of prey availability (Diptera dry biomass). Predictions and their covariates have been back transformed to either the response or unstandardized scale, respectively. Vertical and horizontal hashed lines represent the zero values for both components. Each prediction surface represents a different prey availability group. Prey availability groups are breeding attempts occurring under six different levels of prey availability between 0.0031 g to 0.059 g, representing the minimum to the 99th quantile of observed levels of prey availability. Thus, raw values (points within background) represent the sites scores for the distribution of breeding attempts within each prey availability group. Prediction surfaces of each prey availability group were produced by first calculating the median value of prey availability within each group, delineating out a bounding box of extreme site scores, and then deriving predictions based on these values, all while keeping all other covariates at their mean. The range of colors represent the minimum (deep blue) and maximum (yellow) predicted values of each response variable given site score values.
