## AppendixS2 for "Combined influence of food availability and agricultural intensification on a declining aerial insectivore"

Appendix S2: Supplemental analysis

**Supplemental analysis 1 - Prey availability as the biomass of either Diptera or all insects**

Diptera are the taxa that constitutes a majority of insects found within the diet of Nestling Tree Swallows (Bellavance et al. 2018). In order to validate the assumption that focusing the exploration of prey availability defined as Diptera was valid, we underwent a modeling endeavor in which we compared models treating prey availability as Diptera biomass or the biomass of all insects found within insect traps. It should be noted that during the processing of insect samples, gastropods, large spiders, June bugs (Coleoptera: Melolonthinae), large Lepidoptera, and bumblebees (*Bombus* spp.) were removed, as these are not found in the diet of Tree Swallows. For each response variable we used the full model (i.e., Base + Food * Land for fitness and body condition proxies and Base + Land*Rain + Land*Time for prey availability, see Table 1 and Appendix S1: Table S1 for all covariates found in models). In the following table the AICc values of each of the two models and, when appropriate, the estimate and associated 95% CI of the terms for landcover and prey availability are provided. Data sets were restricted to samples in which the biomass of both all insects and Diptera were available. In each case, models performed better when treating prey availability as Diptera biomass rather than the biomass of all insects.

Table S1: Delta AICc values and coefficient estimates (and their 95% confidence intervals) for identical models fitted to each fitness and body condition proxy and in which the availability of prey was treated as either the biomass of Diptera or the biomass of all insects found within insect traps. Fledging success (N=1,860 broods from 40 farms and 11 years) and prey availability (N=15,225 two-day accumulation of insects) were modeled with GLMMs with a binomial distribution and a logit link function and Gamma distribution with log link function, respectively. The duration of the nestling period (N=1,556 broods), body mass of nestlings (N=6,011), and wing length of nestlings (N=6,065) were all modeled with LMMs. In all cases, the year, farm, and nest box ID were included as random effects with the exception of body mass and wing length as these models also included the brood identification instead of nest box ID.

| Response | Insect | Delta AICc | Prey | Comp.1 | Comp.2 | Comp.1:Comp.2 |
| --- | --- | --- | --- | --- | --- | --- |
| variable | group |  | availability |  |  |  |
| Prey | All insects | 35870.24 |  | 0.00 | 0.01 | -0.01 |
| availability |  |  |  | (-0.03,0.03) | (-0.03,0.06) | (-0.03,0.01) |
|  | Diptera | - |  | **-0.08** | **0.09** | -0.03 |
|  |  |  |  | **(-0.14,-0.03)** | **(0.03,0.15)** | (-0.06,0) |
| Fledging | All insects | 8.71 | 0.00 | 0.01 | **-0.14** | **0.08** |
| success |  |  | (-0.08,0.08) | (-0.07,0.09) | **(-0.28,-0.01)** | **(0.01,0.15)** |
|  | Diptera | - | **0.1** | -0.02 | **-0.14** | **0.08** |
|  |  |  | **(0.02,0.18)** | (-0.1,0.07) | **(-0.27,-0.01)** | **(0.01,0.15)** |
| Duration | All insects | 2.62 | 0.00 | **0.15** | 0.05 | 0.04 |
|  |  |  | (-0.11,0.11) | **(0.06,0.24)** | (-0.11,0.2) | (-0.05,0.12) |
|  | Diptera | - | 0.04 | **0.15** | 0.05 | 0.05 |
|  |  |  | (-0.07,0.16) | **(0.06,0.24)** | (-0.11,0.21) | (-0.03,0.14) |
| Body | All insects | 5.08 | 0.00 | 0.03 | **-0.2** | 0.05 |
| mass |  |  | (-0.11,0.12) | (-0.06,0.13) | **(-0.37,-0.03)** | (-0.04,0.14) |
|  | Diptera | - | 0.08 | 0.02 | **-0.19** | 0.04 |
|  |  |  | (-0.03,0.2) | (-0.08,0.11) | **(-0.35,-0.03)** | (-0.05,0.13) |
| Wing | All insects | 4.28 | 0.03 | **-0.57** | -0.21 | -0.08 |
| length |  |  | (-0.42,0.47) | **(-0.91,-0.22)** | (-0.82,0.4) | (-0.4,0.25) |
|  | Diptera | - | 0.21 | **-0.66** | -0.18 | -0.12 |
|  |  |  | (-0.22,0.64) | **(-1,-0.31)** | (-0.78,0.43) | (-0.45,0.2) |

**Supplemental analysis 2 - Spatio-temporal variation in lower insect groups and larval habitats**

Though outside the scope of this project, we explored the spatio-temporal trends in the biomass of differing insect taxa as a possible hypothesis explaining residual effects of the agricultural intensification gradient on fitness and body condition proxies of breeding Tree Swallows. We investigated this hypothesis using both insects from traps and food boluses delivered to nestlings by their parents. Analyses of the biomass of insect groups was conducted on samples processed as part of a previous publication (full details in Bellavance et al. 2018). Analyses were performed on insect samples from traps from 2011 (N=120) and 2012 (N=35) that coincided with the day, and on the farm, that food boluses were collected. Food boluses were collected throughout the breeding season of 2011 (N=482) and on a subset of 2012 (N=41). It bares repeating that these inferences are derived from data on a small sample from two breeding seasons and a relatively complex study design. Therefore, any conclusions need to take this into account. A main task was to understand the spatio-temporal variation in Diptera of lower taxonomic level in this system. We therefore first separated Diptera from each insect sample or bolus into four principal taxonomic groups (Taxa). These groups included Nematocera, non-Schizophoran Brachycera, Schizophora (Acalyptratae), and Schizophora (Calyptratae). We then used the total biomass of each group from each insect sample or bolus as the response variable. We were further interested in the spatio-temporal variation in biomass of insects either not having an aquatic larval stage (terrestrial) or having an aquatic larval stage (aquatic), henceforth referred to as the source (Twining et al. 2018). We therefore separated the content from each insect sample or bolus into the insects’ respective source, and then used the total biomass of each group from each insect sample or bolus as another response variable. Due to these data being the mass of specific insect groups found within traps or food boluses there were multiple zeros. They were thus analyzed using GLMMs with a tweedie distribution and the Farm ID as a random effect. We further included the Julian date of sample collection, the interaction between Comp.1 and Comp.2, and the percent cover of aquatic habitats within 3 km of the trap or nest box in which the food bolus was collected as fixed effects to both control for and explore their effects (Rioux Paquette et al. 2013, Bellavance et al. 2018, Elgin et al. 2020, Garrett et al. 2021, Berzins et al. 2021). We constructed three models for each combination of response variable (i.e., biomass of trap grouped by Taxa and source and biomass of bolus grouped by Taxa and source). These models focused on interactions between the response variable group (i.e., Taxa or source) and either Comp.1 (Land * Group), Julian date (Julian * Group), or the percent cover of water within 3 km (Water * Group). All covariates were z-transformed prior to analyses. Results presented are the predictions of a single model including only the interaction presented in the figure and the previously stated fixed effects held at their mean. We also provide dot and whisker plots demonstrating the estimate for each coefficient and its associated 95% confidence interval from each model.

We first analyzed the variation in the biomass of major Diptera taxa. We observed that within insect traps, the biomass of Schizophores (both Calyptratae and Acalyptratae) was greater within more agro-intensive landscapes. We did not observe significant variation in the biomass of the primarily aquatic Nematocera and other non-Schizophores with landscape context (Comp.1). Moreover, the percent cover of aquatic habitats within 3 km of traps had a generally negative effect, and yet with a low degree of confidence. The Julian date of collection appeared to have variable effects, dependent on the Diptera group, on biomass (Appendix S2, Figure S1). We further found that in the food boluses delivered to nestlings, the biomass of the primarily aquatic Nematocera slightly decreased with increasing agro intensity and throuhgout the breeding season while increasing with the relative cover of water within 3 km. Moreover, with the exception of Calyptratae, which showed a trend of decreasing with the relative cover of water within 3 km, all other Diptera groups except Nematocera saw similar trends in increasing with agro intensity, the Julian date, and the relative cover of water (Appendix S2, Figure S2).

We next analyzed the variation in the biomass of all identified insect taxa (including non-Dipteran) based on their source (Twining et al. 2018). We observed that within insect traps, a significantly greater proportion of insects have completely terrestrial life-stages. Terrestrial insects decreased in biomass with increasing agro-intensity, Julian date, as well as with the cover of water, and yet with low confidence. However, insects with an aquatic larval stage were not influenced by any of the predictor variables (Appendix S2, Figure S3). Moreover, traps suggested that terrestrial Diptera represent a majority of the insects available, and that their relative availability increases with increasing agro-intensity and Julian date (Appendix S2, Figure S4). Finally, when considering all insects found within food boluses, the provisioning of insects with aquatic life stages decreased with increasing agro-intensity and Julian date (Appendix S2, Figure S5).

The above results highlight that the availability of insects with an aquatic life stage (including Nematocera) were generally less available, and under certain spatio-temporal contexts composed only a (very) small proportion of what was available to food provisioning adult Tree Swallows. The low availability of these insects potentially explains why they are provided at lower rates in general and yet despite this, they are still provided at a greater rate in less agro-intensive conditions. Taken together, these supplemental analyses bring support to the hypothesis that the residual effect of the agricultural gradient on the fitness and body condition proxies studied here may be in part due to variation in dietary quality.

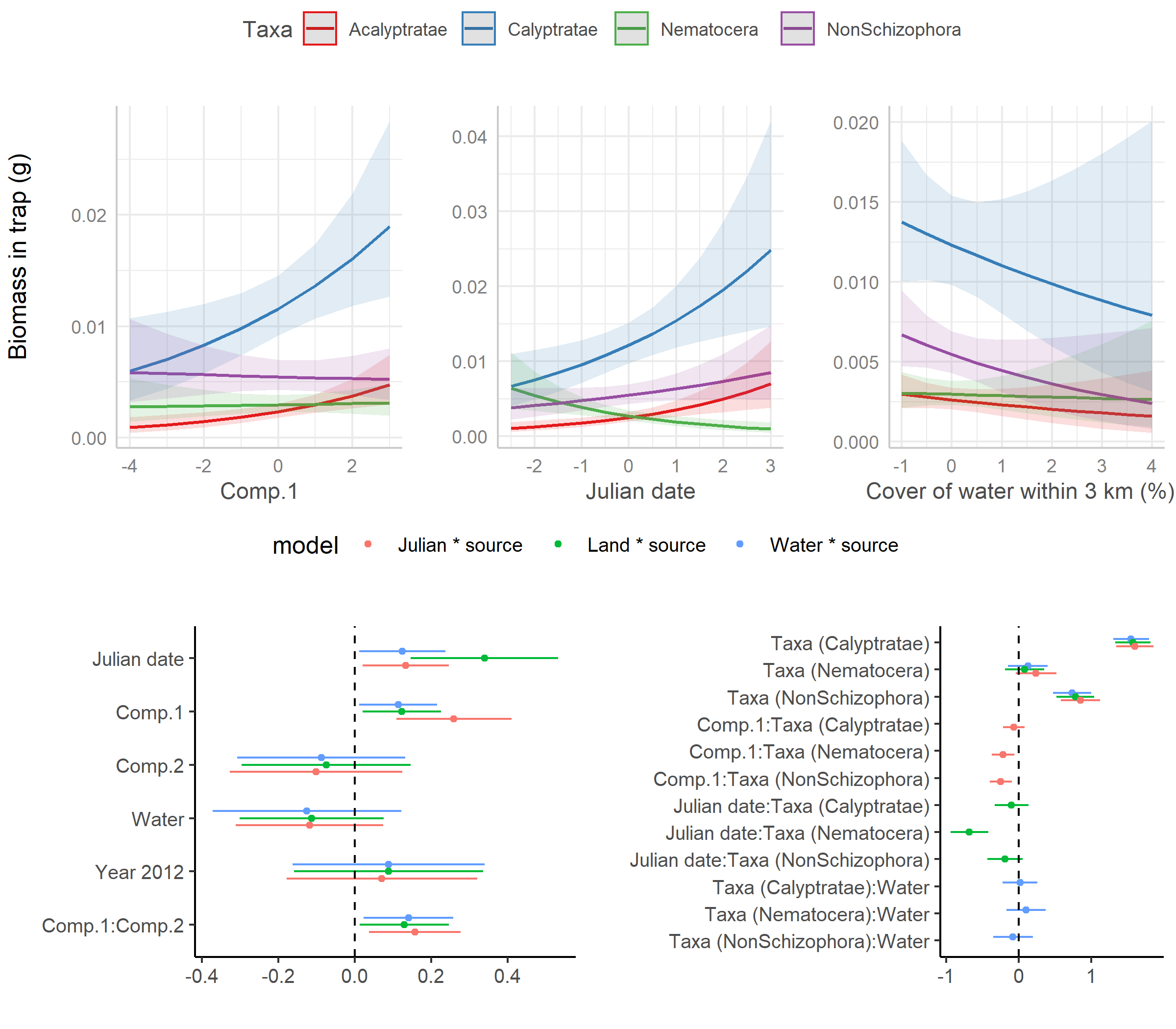

Figure S1: Predicted biomass (per trap) of differing Diptera taxa found within insect traps over two-day sampling periods (N=155) along a gradient of agricultural intensification in southern Quebec, Canada, in 2011 and 2012. Each of the first three plots are predictions from a single model with their 95% confidence intervals plotted against each covariate of interest. In each case, the values of all other covariates have been set to their mean and the predictor under consideration has been back transformed onto its original scale. Second row of plots are the coefficient estimate and its 95% confidence interval for each model term from each of the three models.

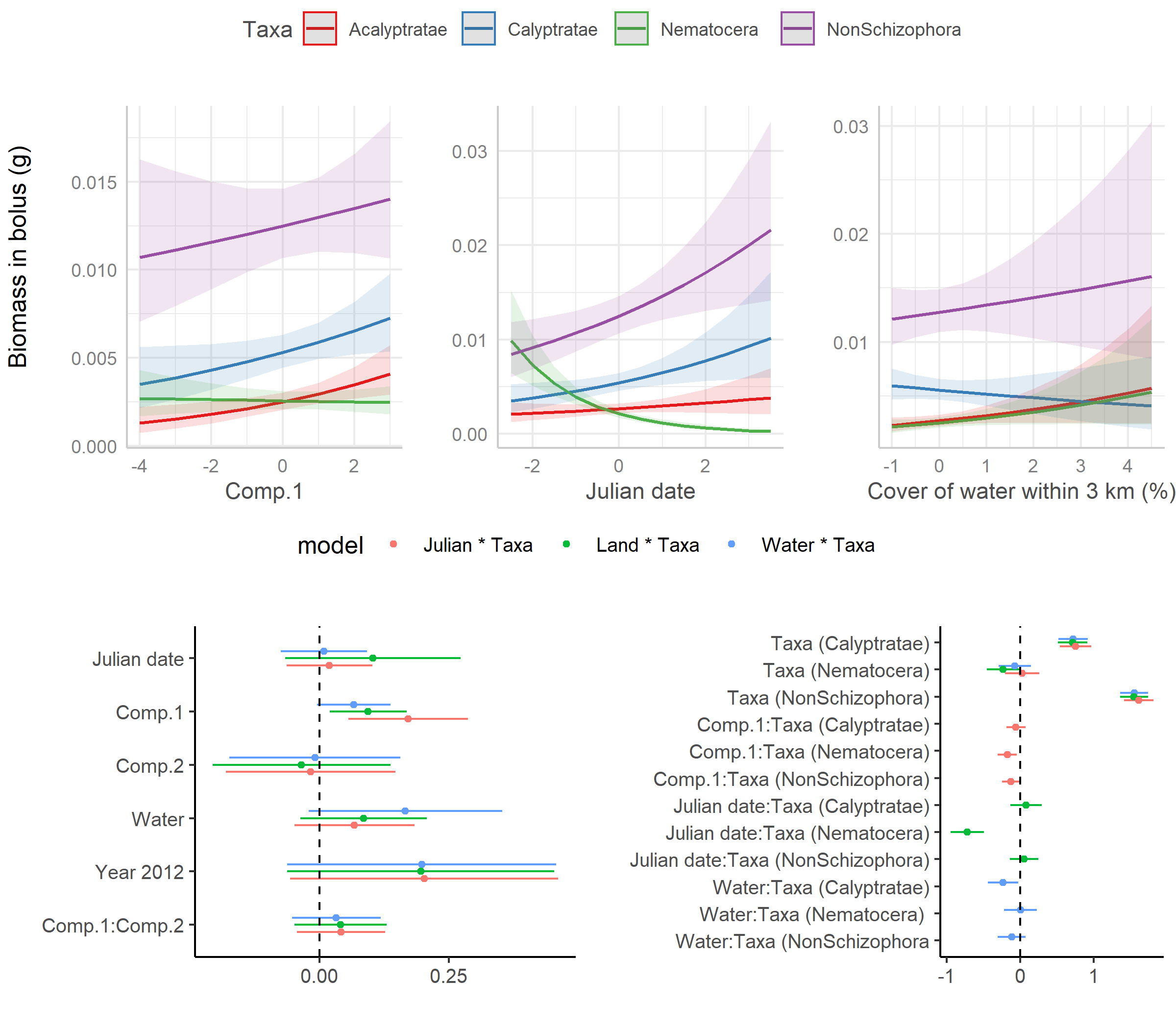

Figure S2: Predicted biomass of differing Diptera taxa found within food boluses (N=523) delivered to Tree Swallow nestlings along a gradient of agricultural intensification in southern Quebec, Canada, in 2011 and 2012. Each of the first three plots are predictions from a single model with their 95% confidence intervals plotted against each covariate of interest. In each case, the values of all other covariates have been set to their mean and the predictor under consideration has been back transformed onto its original scale. Second row of plots are the coefficient estimate and its 95% confidence interval for each model term from each of the three models.

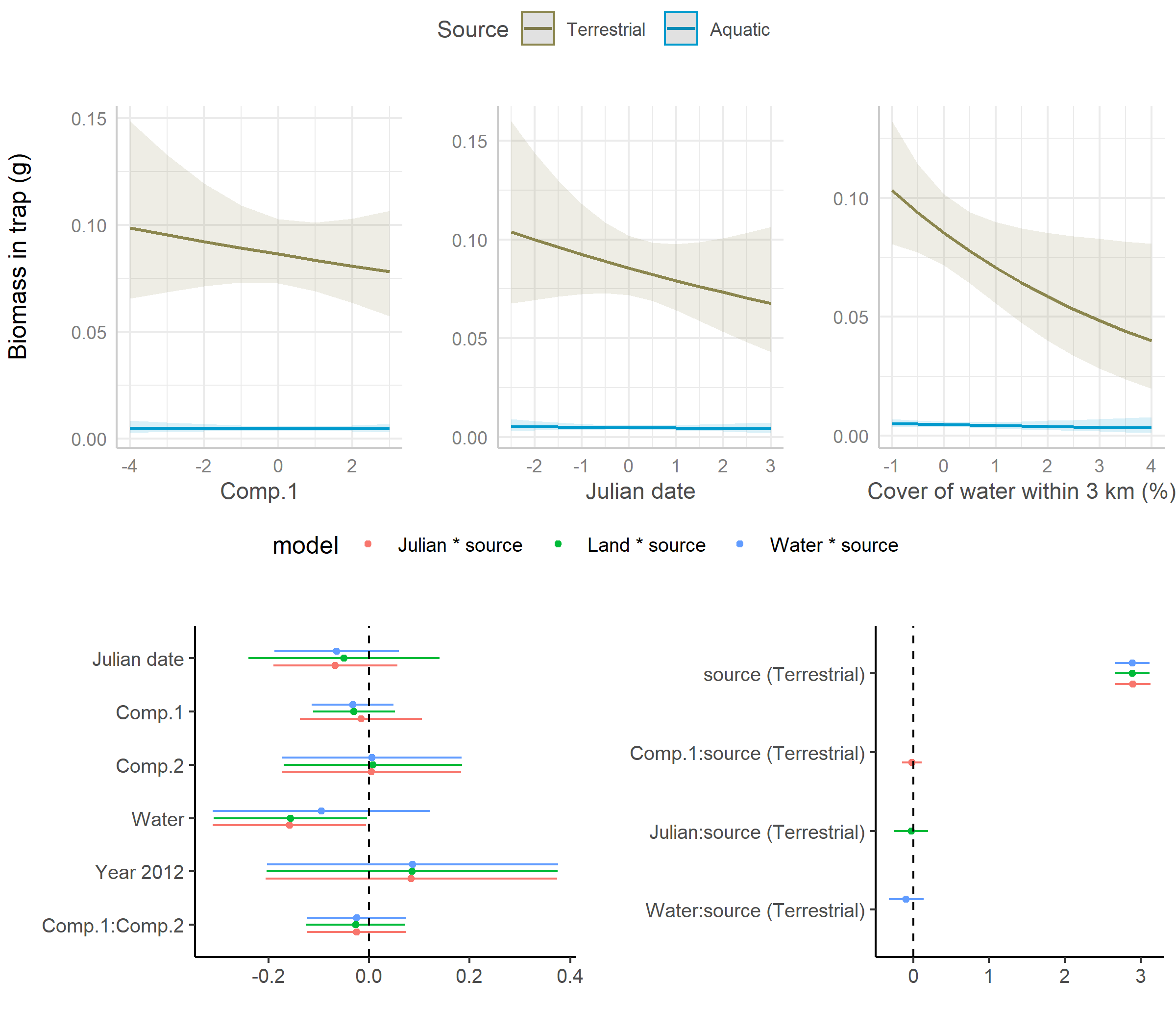

Figure S3: Predicted biomass (per trap) of insects found within insect traps over two-day sampling periods and grouped as whether having or not having an aquatic larval stage (N=155) along a gradient of agricultural intensification in southern Quebec, Canada, in 2011 and 2012. Each of the first three plots are predictions from a single model with their 95% confidence intervals plotted against each covariate of interest. In each case, the values of all other covariates have been set to their mean and the predictor under consideration has been back transformed onto its original scale. Second row of subplots are the coefficient estimate and its 95% confidence interval for each model term from each of the three models.

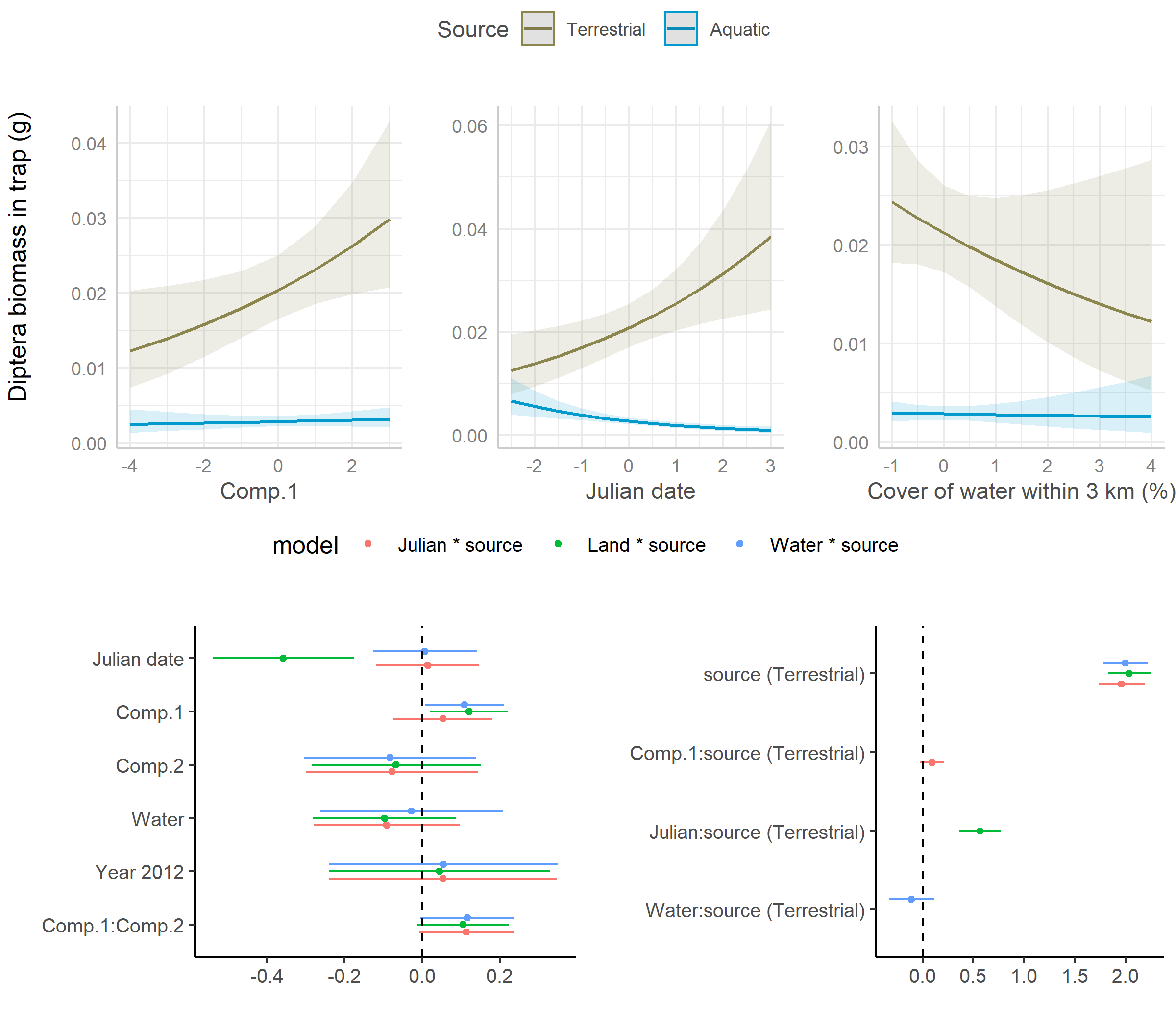
 Figure S4: Predicted biomass (per trap) of Diptera found within insect traps over two-day sampling periods and grouped as whether having or not having an aquatic larval stage (N=155) along a gradient of agricultural intensification in southern Quebec, Canada, in 2011 and 2012. Each of the first three plots are predictions from a single model with their 95% confidence intervals plotted against each covariate of interest. In each case, the values of all other covariates have been set to their mean and the predictor under consideration has been back transformed onto its original scale. Second row of subplots are the coefficient estimate and its 95% confidence interval for each model term from each of the three models.

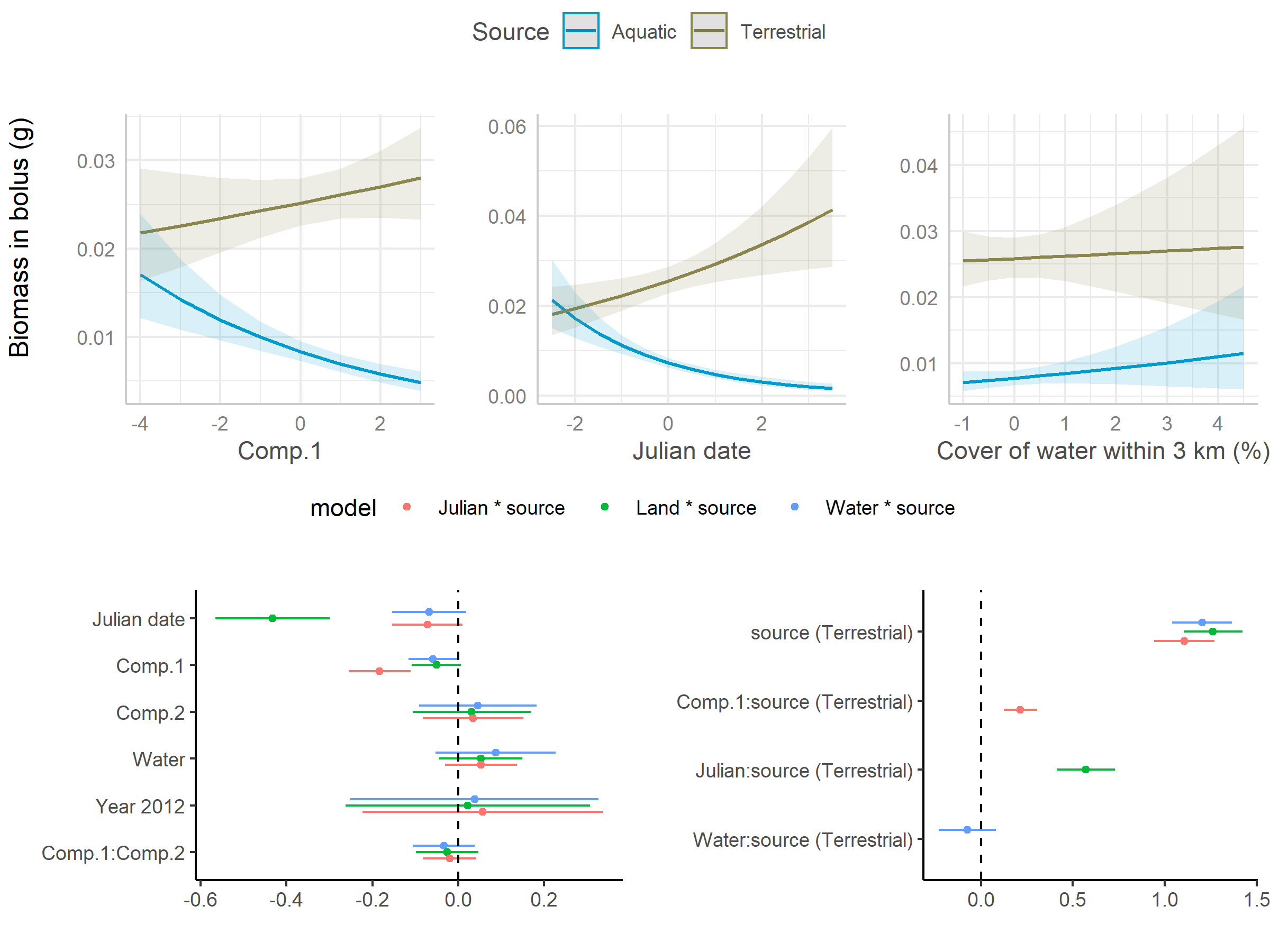

Figure S5 Predicted biomass of insects found within food boluses grouped as whether having or not having an aquatic larval stage (N=523) along a gradient of agricultural intensification in southern Quebec, Canada, in 2011 and 2012. Each of the first three plots are predictions from a single model with their 95% confidence intervals plotted against each covariate of interest. In each case, the values of all other covariates have been set to their mean and the predictor under consideration has been back transformed onto its original scale. Second row of subplots are the coefficient estimate and its 95% confidence interval for each model term from each of the three models.
